# MDA5 is an essential vita-PAMP sensor necessary for host resistance against *Aspergillus fumigatus*

**DOI:** 10.1101/2020.07.06.182154

**Authors:** Xi Wang, Alayna K. Caffrey-Carr, Ko-wei Liu, Vanessa Espinosa, Walburga Croteau, Sourabh Dhingra, Amariliz Rivera, Robert A. Cramer, Joshua J. Obar

## Abstract

RIG-I like receptors (RLR) are cytosolic RNA sensors that signal through the MAVS adaptor to activate interferon responses against viruses. Whether the RLR family has broader effects on host immunity against other pathogen families remains to be fully explored. Herein we demonstrate that MDA5/MAVS signaling was essential for host resistance against pulmonary *Aspergillus fumigatus* challenge through the regulation of antifungal leukocyte responses in mice. Activation of MDA5/MAVS signaling was driven by dsRNA from live *A. fumigatus* serving as a key vitality-sensing pattern-recognition receptor. Interestingly, induction of type I interferons after *A. fumigatus* challenge was only partially dependent on MDA5/MAVS signaling, whereas type III interferon expression was entirely dependent on MDA5/MAVS signaling. Ultimately, type I and III interferon signaling drove the expression of CXCL10. Furthermore, the MDA5/MAVS-dependent interferon response was critical for the induction of optimal antifungal neutrophil killing of *A. fumigatus* spores. In conclusion, our data broaden the role of the RLR family to include a role in regulating antifungal immunity against *A. fumigatus*.

**KEY POINTS:**

- MDA5 is essential for maintaining host resistance against Aspergillus fumigatus
- MDA5 serves as a critical vitality sensor after fungal challenge
- MDA5 is essential for IFNλ expression and anti-fungal neutrophil killing

## INTRODUCTION

*Aspergillus fumigatus* is a ubiquitous environmental mold that humans inhale on a daily basis. Multiple environmental surveys have demonstrated a range of several hundred to thousands of *A. fumigatus* conidia can be inhaled daily (1–3). Individuals with normal immune systems readily clear *A. fumigatus* conidia from their airways without problems. However, immunocompromised individuals are at significantly greater risk of developing invasive pulmonary aspergillosis (IPA)^4^. IPA is a life-threatening disease with over 300,000 cases globally and a high mortality rate of 30% - 50% (4). Patients at increased risk of developing IPA include those receiving chemotherapy treatments for cancer and patients receiving immunosuppressive regimens to prevent graft-versus-host disease (GVHD) following hematopoietic stem cell transplants or solid organ transplants (5–9). Furthermore, epidemiology studies predict a steady increase of IPA cases in the future due to the projected continual increase of immune compromised patients (10). Additionally, individuals with single nucleotide polymorphisms (SNPs) or primary immunodeficiencies in key antifungal signaling or effector pathways, such as CARD9 (11), pentraxin 3 (12, 13), or NADPH oxidase (14, 15) are highly susceptible to IPA. Thus, these clinical observations demonstrate that host innate immune responses are imperative in deactivating viable fungal conidia to avoid fungal establishment in the lung, and failure of such processes result in fungal germination and fungal growth and dissemination, ultimately culminating in IPA.

If the physical barriers of the lungs are bypassed by the inhaled *A. fumigatus* conidia, airway epithelial cells and lung-resident macrophages comprise the first line of defense against inhaled conidia. It is well established that lung resident macrophages, including alveolar macrophages (16, 17) and CCR2^+^ monocytes (18, 19) are critical for initiating the early inflammatory milieu necessary for the recruitment of innate immune cells to the respiratory tract to mediate host resistance against invasive disease. These innate immune effector cells include neutrophils (20, 21), inflammatory monocytes (18, 19), NKT cells (22), and plasmacytoid dendritic cells (23–25). The cytokine/chemokine signaling networks necessary to regulate the recruitment of inflammatory cells to the respiratory tract are rapidly emerging, but still much remains to be learnt.

Recently, it has become well appreciated that the host inflammatory response induced by *A. fumigatus* is tightly tuned to the virulence of the individual strain under study (26–30). Being able to recognize pathogenic traits of microbes is key to appropriately responding to infection (31), one such trait is being able to resist killing and remain viable (32). Two crucial inflammatory pathways in signaling bacterial vitality are inflammasome-dependent secretion of IL-1β and induction of the interferon response, which both can be driven by bacterial RNA (33, 34). We and others have previously demonstrated that inflammasomes are essential for IL-1β production after *A. fumigatus* challenge (35–38). Interestingly, Kanneganti and colleagues have shown to the NLRP3 and AIM2 inflammasomes could be activated by fungal RNA and DNA, respectively (35). Moreover, both type I and type III interferons have been shown to be essential for host resistance against pulmonary *A. fumigatus* challenge (39), but the fungal PAMP and host pattern-recognition pathways leading to type I and type III interferon expression following *A. fumigatus* challenge have not been well elucidated.

Type I and type III interferon responses are best understood in the context of virus infections. Following virus infections, Toll-like receptors (including TLR3, TLR7/8, TLR9), cGAS/STING, and RIG-I-like receptors (RIG-I and MDA5) are known to be critical in the induction of type I and type III interferons (review by (40–42)). However, during fungal infections much less is currently known. Both TLR3 and TLR9 are known to be involved in the antifungal defenses against *A. fumigatus* (43–46), but only their role in the induction of type I interferons has been explored (45, 47). Following *Candida albicans* infection TLR7 is known to be involved in the induction of type I interferons (48, 49). Moreover, a Dectin-1/Syk/Irf5 signaling network has been shown to be critical in the induction of type I interferons following *C. albicans* challenge (50). Finally, Dectin-1 has been shown to be partially involved in the induction of type I and III interferons following respiratory *A. fumigatus* challenge (51). However, the role of cytosolic sensing pattern-recognition receptors systems in the antifungal interferon response has not been well studied.

Our data demonstrate that MDA5/MAVS-dependent signaling is necessary for host resistance against *A. fumigatus* infection through the accumulation and activation of antifungal effector functions of neutrophils in the fungal challenged airways. MAVS signaling drove expression of both type I and type III interferon, which ultimately drove the expression of the interferon inducible chemokines CXCL9 and CXCL10. Overall, our study reveals a critical role for MDA5 in host anti-fungal immunity, which supports a broader role of MDA5 in monitoring the health of the host cytosol.

## MATERIALS AND METHODS

### Mice

C57BL/6J (Jackson Laboratory, Stock #000664), *Mavs^-/-^* (Jackson Laboratory, Stock #008634), and *Ifih1^-/-^* (Jackson Laboratory, Stock # 015812) were all originally purchased from Jackson Laboratories and sequentially bred in-house at Geisel School of Medicine at Dartmouth. *Ifnar^-/-^*, *Ifnlr^-/-^*, and *Ifnar^-/-^ Ifnlr1^-/-^* DKO were kindly provided by Dr. Sergei Kotenko and Dr. Joan Durbin (52) and bred in-house at Rutgers – New Jersey Medical School. Wild-type B6.129F2 (Jackson Laboratory, Stock #101045) mice, used as controls for the *Mavs^-/-^* mice, were purchased directly from Jackson Laboratories. All mice were 8-16 weeks of age at the time of challenge. Both male and female mice were used for these experiments. All animal experiments were approved by either the Rutgers – New Jersey Medical School Institutional Animal Care and Use Committee or Dartmouth College Institutional Animal Care and Use Committee.

### Preparation of Aspergillus fumigatus conidia

*A. fumigatus* CEA10 and Af293 strains were used for this study. Each strain was grown on glucose minimal media (GMM) agar plates for 3 days at 37°C. Conidia were harvested by adding 0.01% Tween 80 to plates and gently scraping conidia from the plates using a cell scraper. Conidia were then filtered through sterile Miracloth, were washed, and resuspended in phosphate buffered saline (PBS), and counted on a hemocytometer.

### Preparation of swollen conidia and heat-killing

*A. fumigatus* CEA10 conidia were collected, as described above. To generate a homogenous population of swollen conidia, conidia were resuspended at 5×10^6^ conidia per ml in RPMI containing 0.5 μg/ml of voriconazole (Sigma-Aldrich) and incubated at 37°C for 15 h, as previously done (53). Conidia were then washed twice and adjusted to 4×10^8^ conidia per ml in PBS-Tween 80. To heat-kill the swollen conidia, the conidial suspension was incubated in a 100°C water bath for 30 minutes. The conidial concentration and efficiency of heat-killing was verified by plating on GMM agar.

### Aspergillus fumigatus pulmonary challenge model

Mice were challenged with *A. fumigatus* conidia by the intratracheal (i.t.) route. Mice were anesthetized by inhalation of isoflurane; subsequently, mice were challenged i.t. with ∼4 × 10^7^ *A. fumigatus* conidia in a volume of 100 µl PBS. At the indicated time after *A. fumigatus* challenge, mice were euthanized using a lethal overdose of pentobarbital. Bronchoalveolar lavage fluid (BALF) was collected by washing the lungs with 2 ml of PBS containing 0.05*M* EDTA. BALF was clarified by centrifugation and stored at -20°C until analysis. After centrifugation, the cellular component of the BALF was resuspended in 200 μl of PBS and total BAL cells were determined by hemocytometer count. BALF cells were subsequently spun onto glass slides using a Cytospin4 cytocentrifuge (Thermo Scientific) and stained with Diff-Quik (Siemens) or Hema 3™ Stat Pack (Fisher Scientific) stain sets for differential counting. For histological analysis lungs were filled with and stored in 10% buffered formalin phosphate for at least 24 hours. Lungs were then embedded in paraffin and sectioned into 5-micron sections. Sections were stained with Grocott-Gomori methenamine silver (GMS) using standard histological techniques to assess lung inflammatory infiltrates and fungal germination, respectively. Representative pictures of lung sections were taken using either the 20X or 40X objectives on an Olympus IX73 with a Zeiss Aziocam 208 color camera.

### Luminex assay for cytokine and chemokine secretion from experimental murine models of invasive aspergillosis

BALF and lung homogenates from B6/129F2 mice and *Mavs^-/-^* mice challenged with *A. fumigatus* 12 or 48 h prior were analyzed for cytokines and chemokines using Milliplex Mouse Cytokine & Chemokine 32-plex (Millipore). Plates were read using a BioPlex 200 (Bio-Rad) in the Immune Monitoring and Flow Cytometry Core Facility at Dartmouth College.

### Quantitative RT-PCR analysis

Total RNA from lungs was extracted with Trizol (Invitrogen). One microgram of total RNA was reverse transcribed using High Capacity cDNA Reverse Transcription Kit (Applied Biosystems). TaqMan Fast Universal Master Mix (2X) No AmpErase UNG and TaqMan probes (Applied Biosystems, Catalog #4331182) for *Ifna2* (Mm00833961_s1), *Ifnb1* (Mm00439552_s1), *Ifnl2/3* (Mm0404158_gH), *Ifng* (Mm01168134_m1), *Cxcl9* (Mm00434946_m1), and *Cxcl10* (Mm00445235_m1) were used and normalized to *Gapdh* (Mm99999915_g1). Gene expression was calculated using ΔΔCT method relative to naïve sample.

### Western blot analysis

Total protein from whole lungs was extracted using RIPA buffer. Total protein was quantified using a DC™ Protein Assay (Bio-Rad). For Western Blot analysis, 40 μg of total protein was loaded per well. Gels were run at 100V for ∼1 hour at room temperature and then transferred to PVDF membranes at 100V for ∼1 hour at 4°C. After transfer, the membranes were blocked with TBST + 5% milk for 1 hour at room temperature. Following blocking, blots were rinsed twice with TBST buffer. Staining with primary antibodies was done in TBST with 5% BSA at 4°C for overnight on a roller. Primary antibodies used for these studies were rabbit anti-STAT1 (Cell Signaling, #9172S, 1/1000), rabbit anti-pSTAT1 (Cell Signaling, #7649P, 1/1000 dilution), and rabbit anti-β-actin (Abcam, ab75186, 1/40,000). After primary antibody staining blots were washed four times with TBST. Blots were then stained with a donkey anti-rabbit IgG conjugated to HRP (Cell Signaling, #7074P2, 1/4000 dilution) in TBST + 5% milk for 1 hour at room temperature with gentle shaking. After secondary antibody staining blots were washed four times with TBST. Blots were developed using ECL Clarity (Bio-Rad) for 5 minutes, then analyzed using an Alpha Innotech FluorChem Q imager.

### Generation of a Fluorescent Aspergillus fumigatus reporter (FLARE) strain in the CEA10 background

CEA10:H2A.X*^A. nidulans^ ptrA* was constructed in two steps. First, *gpdA*(p) was amplified from DNA isolated from strain *A*. *nidulans* A4 (Source: FGSC) using primers RAC2888 and RAC2799. Histone variant H2A.X (AN3468) was amplified (without stop codon) from A4 DNA using primers RAC4582 and RAC4583. mRFP fragment was amplified from plasmid pXDRFP4 (Source: FGSC) using primers RAC2600 and RAC4575 and terminator for *A*. *nidulans trpC* gene was amplified using primers RAC2536 and RAC2537. The four fragments were then fused together with primers RAC1981 and RAC4134 using fusion PCR resulting in H2A.X first round fragment as described earlier (54). All primers are listed in Supplemental Table 1.

Secondly, we targeted integration of the H2A.X rfp to the intergenic locus between AFUB_023460 and AFUB_023470. For this, left homology arm was amplified from CEA10 genomic DNA using primers RAC3873 and RAC3874. Right homology arm was amplified with primers RAC3875 and RAC3876. Dominant selection marker gene *ptrA* conferring resistance to pyrithiamine hydrobromide was amplified from plasmid pSD51.1 with primers RAC2055 and RAC2056. The four fragments were then fused together with primers RAC3877 and RAC3878 using fusion PCR as described earlier (54). After the construct generation, polyethylene glycol mediated transformation of protoplast was performed as described earlier (54). Transformants were screened by PCR (data not shown) and confirmed with southern blot analysis as described earlier (55). mRFP fluorescence was confirmed with FACS (Fluorescence activated cell sorting) analysis.

### Fluorescent Aspergillus fumigatus reporter (FLARE) assay

To measure both conidial uptake and viability in distinct immune cell populations we used fluorescent *Aspergillus* reporter (FLARE). The CEA10-based FLARE was labelled with AlexaFlour 633 membrane labeling, as described elsewhere (56). Bronchoalveolar lavage (BAL) and lung cell suspensions were prepared as described elsewhere (56) and stained with anti-Ly6G (1A8), anti-CD11b (M1/70), anti-CD11c (HL3), anti-CD45 (30-F11), anti–CD206 (C068C2) and a fixable viability dye (eBioscience; Catalog #65-0865-14). Neutrophils were identified as CD45^+^CD11b^+^Ly6G^+^, alveolar macrophages as CD45^+^CD11b^+^Ly6G^−^CD206^dim^, interstitial macrophages as CD45^+^CD11b^+^Ly6G^−^CD206^−^, and monocytes as CD45^+^CD11b^dim^Ly6G^−^CD206^+^ cells, as previously defined (57, 58). Flow cytometric data were collected on a CytoFLEX (Beckman Coulter). All data was analyzed using FlowJo software.

### Primary murine fibroblast cultures

Ears from 6-12 week-old C57BL/6J and *Ifih1^-/-^* mice were collected, washed in 70% EtOH, then PBS, and finally minced using scissors. For each ear, 500 μl of 1000 U/ml collagenase (Gibco, REF 17101-015) in HBSS was used to digest them at 37°C for 25 minutes. The tissues were then centrifuged at 500 *x g* for 3 minutes and then washed once with HBSS. Ears were subsequently digested with 500 μ 20 minutes in 37°C. The tissues were then centrifuged again at 500 *x g* for 3 minutes, and the trypsin was discarded and replaced by 0.5ml fibroblast media [10% FCS, 1% MEM Non-essential amino acids, 1% pencillin-streptomycin in DMEM]. The tissues were then pushed through a sterile 40 μm cell strainer via the plunger of a 3ml syringe to obtain single cell suspensions. For every ear, 25ml of fibroblast media was used to initiate the murine fibroblast culture in tissue culture flasks and a media change was conducted every two days. The fibroblasts were usually confluent and harvested on day 6 by 0.25% trypsin digest for 5min in 37°C and then collected with fresh fibroblast media.

### Isolation of total RNA from Aspergillus fumigatus

*A. fumigatus* conidia was harvested from GMM plates and inoculated at 1 x 10^6^ per ml in liquid GMM. Cultures were grown for 24h in 37°C with shaking at 250 rpm in an orbital shaker. Mycelia were then filtered and wrapped in sterile Miracloth, dried by repeated paper towel absorption, and weighed. For every 50 mg of mycelium, 1 ml of Trisure (Bioline) was added and mixed. The Trisure-mycelium mixture was frozen by liquid nitrogen to release cellular content and then homogenized using a mortar and pestle. For every 1 ml of Trisure used, 0.2 ml chloroform was added and vortexed, and the solution was centrifuged for 15 min at 12000 *x g* at 4°C. The aqueous phase was then transferred to a new tube and 0.5 ml of isopropanol was added, vortexed, and centrifuged for 10 min at 12000 *x g* at 4°C. The supernatant was then removed, and the pellet was washed by 1 ml 75% ethanol via vortexing followed with centrifuging for 5 min at 7500 *x g* at 4°C. The 75% ethanol was then removed to allow a complete air dry of the RNA, followed by dissolving in molecular biology grade water and stored in -80°C.

### Stimulation of murine fibroblasts with fungal RNA

Freshly harvested murine fibroblasts were dosed at 5 x10^4^/ml in fibroblast media and transferred to 24-well tissue culture treated plates with 0.5 ml per well. The plate was incubated for one day to allow the attachment of murine fibroblasts. Fibroblasts were then transfected via LyoVec™ (Invivogen)/RNA complexes. For every 200 μl of LyoVec™, 2 μg of either poly-IC (Invivogen), 5’-ppp RNA (Invivogen), or *A. fumigatus* RNA was diluted in 80 μl molecular biology grade water, mixed, and incubated in room temperature for 30 minutes to assemble the liposome. The assembled liposome complexes were then added to 5 ml of fibroblast medium and 0.5 ml of the above transfection media was transferred to each well of 24-well plates after removal of the original medium. Twenty-four hours after transfection the supernatant was collected for subsequent ELISA analysis.

### ELISA analysis for cytokine and chemokine secretion

Cell culture supernatants from fibroblast stimulated with *A. fumigatus* RNA were analyzed by ELISA for CXCL10 (R&D Systems, Cat. DY466) and Interferon alpha (Invitrogen, Cat. BMS6027TWO). Plates were read using an Epoch BioTek Gen5 microplate reader at 450nm, and the background was subtracted at 570nm.

### Statistical analysis

Statistical significance for *in vitro* and *ex vivo* data was determined by a Mann-Whitney U test, one-way ANOVA using a Tukey’s or Dunn’s post-test, or two-way ANOVA with Tukey’s post-test through the GraphPad Prism 7 software as outlined in the figure legends. Mouse survival data were analyzed with the Mantel-Cox log rank test using GraphPad Prism.

## RESULTS

### Induction of an optimal interferon response by *Aspergillus fumigatus* is dependent on viable conidia

The polysaccharide-rich cell wall is a major driver of the inflammatory response against *A. fumigatus* and other fungi. Following *C. albicans* challenge the Dectin-1 (*Clec7a*) receptor is critical in the induction of type I interferons following (50). Dectin-1 is only partially responsible for induction of the type I and type III interferon response induced by *A. fumigatus* (51). To test whether the fungal cell wall was essential for driving the type I and type III interferon response, we stimulated C57BL/6J mice with 4×10^7^ live or heat-killed swollen conidia of the CEA10 strain of *A. fumigatus*. Forty hours post-challenge with live swollen conidia there was an induction of IL-28 (IFN-λ), CXCL10, and TNFα secretion (Figure 1). In contrast, when challenged with heat killed swollen conidia secretion of both IL-28 (IFN-λ) and CXCL10 was markedly reduced, while TNFα was still induced (Figure 1). These finding demonstrate that while cell carbohydrates in heat-killed swollen conidia still drove robust inflammatory cytokines (such as TNF α), as has previously been demonstrated (53, 59), there is an alternative pattern-recognition receptor pathway which is largely responsible for the interferon response induced by live *A. fumigatus* conidia.

**Figure 1.**
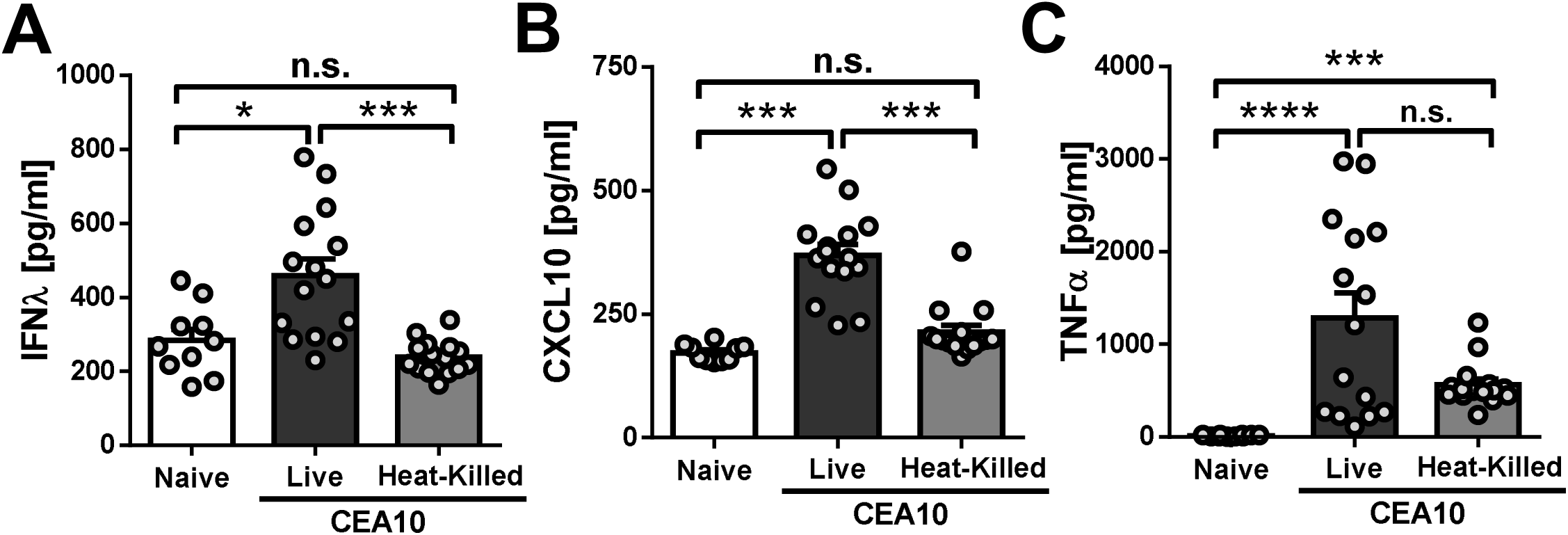
**Heat-killed swollen conidia have a decreased ability to induce CXCL10.** C57BL/6J mice were challenged i.t. with 4×10^7^ CEA10 live or heat-killed swollen conidia. Forty hours later BALF was collected and IL-28/IFN-α **(A)**, CXCL10 **(B)**, and TNF **(C)** levels were determined by ELISA. Data are pooled from 2 independent experiments with 10-15 total mice per group. Statistical significance was determined using a one-way ANOVA with Dunn’s post-test (*p<0.05; **p < 0.01).

### dsRNA from *Aspergillus fumigatus* drives an MDA5-dependent interferon response

Vitality sensing during bacterial infection through recognition of bacterial RNA has been shown to be critical for the induction of protective immune responses through the secretion of both IL-1β and type I interferons (33). Thus, we next wanted to test whether *A. fumigatus* RNA could drive interferon release. To test this hypothesis, primary murine fibroblasts from C57BL/6J mice were stimulated with naked or liposome packaged total RNA from the Af293 strain of *A. fumigatus.* As expected, untreated fibroblast or empty liposome treated did not drive significant secretion of IFNα or CXCL10, while liposome packaged polyI:C drove significant secretion of both (Figure 2A). Liposome-packaged Af293 RNA drove secretion of both IFNα and CXCL10, while naked Af293 could not drive their production (Figure 2A). Taken together, these data demonstrate that intracellular sensing of *A. fumigatus* RNA can drive an interferon-dependent inflammatory response.

**Figure 2.**
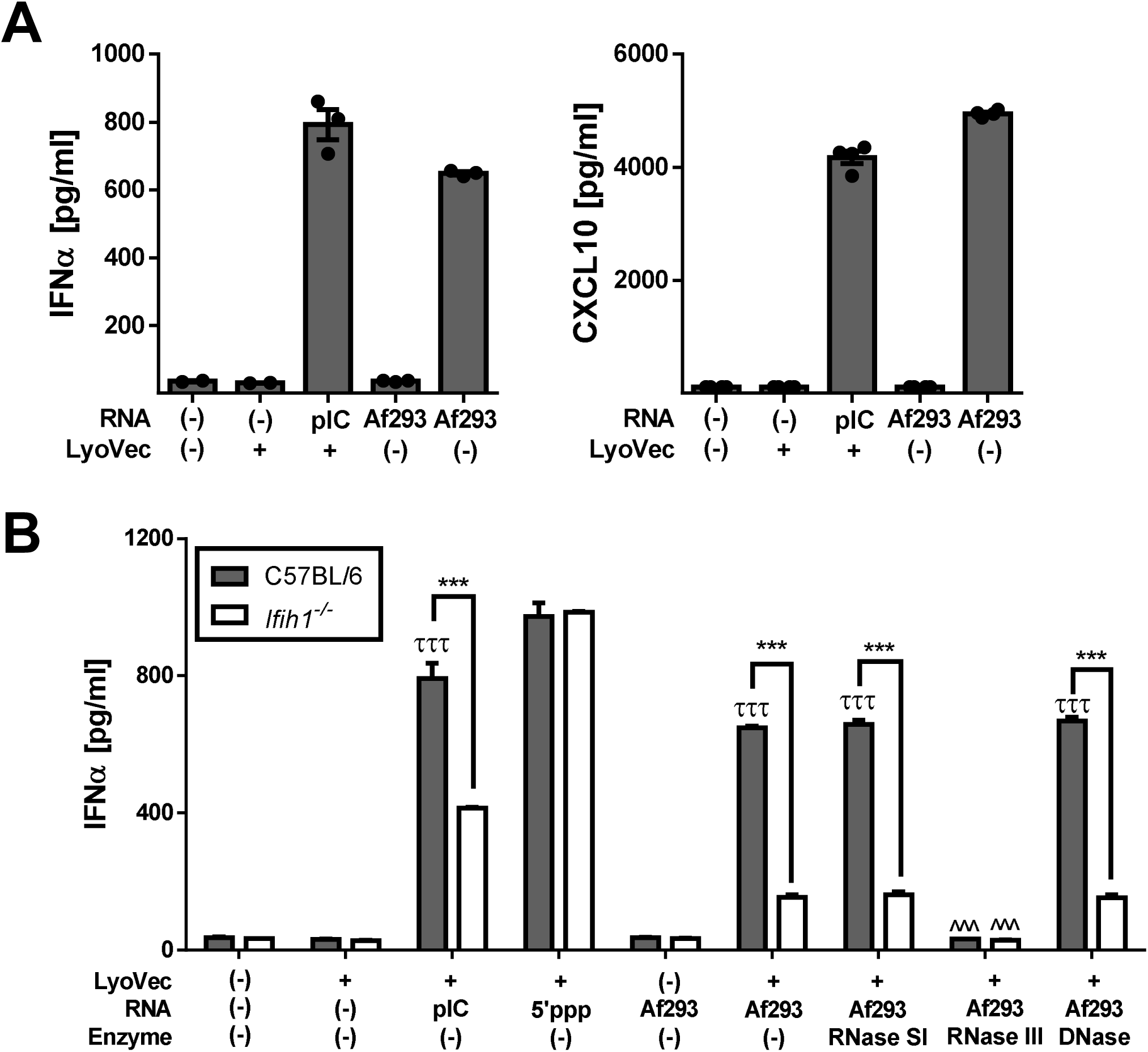
***Aspergillus fumigatus* RNA induces an interferon response in a partially MDA5-dependent manner. (A)** Primary murine fibroblasts from C57BL/6J mice were stimulated with LyoVec™ encapsulated polyI:C (pIC) or total RNA isolated from Af293 for 18 h. After stimulation cell supernatants were collected and analyzed for IFN α (left) and CXCL10 (right) by ELISA. Data from 2 independent experiments with 3 samples per group. **(B)** Primary murine fibroblasts from C57BL/6J and *Ifih1^-/-^* mice were stimulated with LyoVec™ encapsulated polyI:C (pIC) (MDA5 agonist), 5’-ppp (RIG-I agonist), or total RNA isolated from Af293 for 18 h. The total RNA pool isolated from Af293 was also treated with either RNase S1, RNase III, or DNase to degrade ssRNA, dsRNA, or DNA, respectively prior to encapsulation in LyoVec™. After stimulation cell supernatants were collected and analyzed for IFNα by ELISA. Data from 2 independent experiments with 3 samples per group. Statistical significance was determined by a two-way ANOVA with a Tukey’s post-test (τττ p<0.001 – LyoVec™ only vs. experimental group; *** p<0.001 – B6 vs. *Ifih1^-/-^*; ^^^ p<0.001 – enzyme treated vs. Af293).

Intracellular sensing of RNA is mediated by the RIG-I-like receptor family, composed of RIG-I and MDA5, for the induction of interferons. Both RIG-I and MDA5 recognize distinct structures in foreign RNA (60, 61). To test whether ssRNA or dsRNA from *A. fumigatus* was required to drive type I IFN secretion, we extended these *in vitro* fibroblast stimulation studies by pre-treating the *A. fumigatus* RNA pool with either RNase S1 or RNase III, which will selectively cleave ssRNA and dsRNA, respectively, prior to liposome packaging. We also treated the fungal RNA pool with DNase I as a control. Similar to our previous result, total *A. fumigatus* RNA packaged in liposomes was able to induce IFNα secretion from murine fibroblast (Figure 2B). Pre-treatment of the *A. fumigatus* RNA with either RNase S1 or DNase I had no impact on the secretion of IFNα (Figure 2B). In contrast, pre-treatment of the *A. fumigatus* RNA with RNase III completely ablated the secretion of IFNα from primary murine fibroblasts (Figure 2B). These data suggest that recognition of double-stranded fungal RNA in the cytosol is necessary for the interferon response induced by *A. fumigatus*.

To date the most well characterized intracellular receptor for dsRNA is MDA5 (60). To test whether *A. fumigatus* RNA initiates an interferon response in a MDA5-dependent manner we treated primary murine fibroblasts isolated from C57BL/6J or *Ifih1^-/-^* mice liposome packaged fungal RNA. As positive controls, we utilized high molecular weight polyI:C, a known MDA5 ligand, and 5’ppp-RNA, a known RIG-I ligand. Eighteen hours are stimulation cell culture supernatants were collected for ELISA analysis to assess IFNα secretion. Unstimulated or LyoVec™ only stimulated fibroblast did not secrete any IFNα (Figure 2B). As positive controls, 5’ppp-RNA stimulated equivalent secretion of IFN α from both wild-type and *Ifih1*-deficient fibroblasts; while high molecular weight polyI:C stimulated robust IFN α secretion from wild-type fibroblast, which was significantly reduced in *Ifih1*-deficient fibroblasts (Figure 2B). The induction of IFNα secretion by *A. fumigatus* RNA was partially dependent on MDA5, like what we also observed with high molecular weight polyI:C (Figure 2B). Interestingly, RNase III treatment ablated both the MDA5-dependent and -independent IFN-α response (Figure 2B), which suggests there is another dsRNA receptor that contributions to the IFN-α response in response to encapsulated fungal RNA.

### MAVS-dependent signaling is required for interferon expression after *Aspergillus fumigatus* challenge

During viral infections RLR family members are critical in initiating both the type I and type III interferon responses (62). Moreover, stimulation of *Ifih1-*deficient murine macrophages with *C. albicans* resulted in a slight decrease in *Ifnb* mRNA expression, which did not reach significance (63). However, the specific role of MDA5/MAVS in initiating the induction of a broad range of interferons to other fungal pathogens has not been explored. To test the role of MAVS signaling in the induction of interferons after respiratory challenge with *A. fumigatus*, we challenged B6.129F2 mice and *Mavs^-/-^* mice with 4×10^7^ conidia of CEA10. We collected lungs from the *A. fumigatus* challenged mice at 3 and 48 h post-inoculation for quantitative RT-PCR analysis to assess interferon expression patterns. Similar to previous observations with C57BL/6J mice (39), wild-type B6.129F2 mice expressed high levels of type I interferons, *Ifna4* (Figure 3A) and *Ifnb* (Figure 3B), at 3 h post-inoculation that waned with time. Both type II interferon (*Ifng*) (Figure 3D) and type III interferon (*Ifnl2/3*) (Figure 3C) were expressed at higher levels at 48 h post-inoculation in wild-type B6.129F2 mice. Interestingly, in the *Mavs-*deficient mice type I interferon levels were only decreased by approximately 50% (Figure 3A-B), whereas the expression of type II and type III interferons were nearly completely ablated in the absence of MAVS (Figure 3C-D). Thus, MAVS-dependent signaling is essential for the late expression of type III interferon and only partially responsible for early expression of type I interferons.

**Figure 3.**
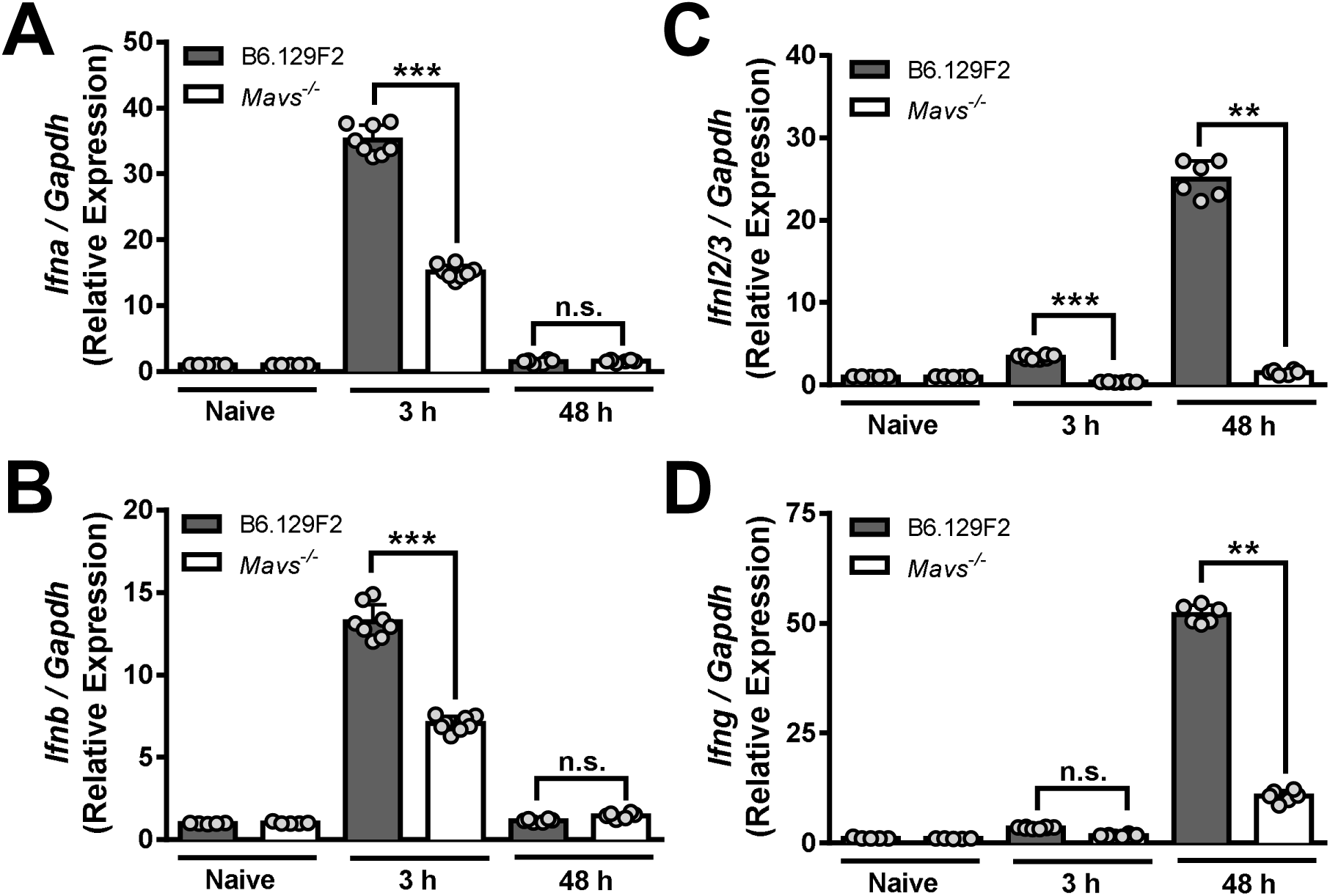
***Mavs* mice have decreased interferon mRNA levels after *Aspergillus fumigatus* challenge particularly at later times.** B6.129F2 and *Mavs* mice were challenged with 4×10 conidia of CEA10. Lungs were collected 3 or 48h post-inoculation. Total RNA was extracted from whole lungs. Gene expression as determined by quantitative reverse transcription polymerase chain reaction (qRT-PCR) using TaqMan probes for *Ifna* **(A)**, *Ifnb* **(B)**, *Ifnl2/3* **(C)**, and *Ifng* **(D)**, which were normalized to *Gapdh* expression. Bars represent data means ± SEM with each dot representing individual mice. Data are representative of results from at least 2 independent experiments with at least 3 mice per group. Data were analyzed using a Mann-Whitney U-test (** p < 0.01; *** p < 0.0001).

To confirm that the expression of interferons resulted in down-stream IFN signaling, we next examined the phosphorylation of STAT1. STAT1 phosphorylation is essential for the signaling of all interferon classes. Lungs from C57BL/6J mice challenged 6, 24, or 48 h prior with either the CEA10 or Af293 strains were collected, and total proteins were extracted for Western blot analysis. Both the CEA10 and Af293 strains induced a bi-phasic induction of STAT1 phosphorylation (Figure 4A). Specifically, significant STAT1 phosphorylation was observed at 6 and 48 h after challenge, but minimal phosphorylation was observed at 24 h (Figure 4A). These data fit with our earlier observation that type I interferons are produced by 3 h post-challenge, while type II and type III interferons are highly expressed at later times (Figure 3). We next examined the phosphorylation of STAT1 in the lungs of B6.129F2 and *Mavs^-/-^* mice challenged with the CEA10 stain of *A. fumigatus* at 6 and 48 h after challenge, times when we saw high levels of STAT1 phosphorylation. Like C57BL/6J mice, wild-type B6.129F2 mice induced robust phosphorylation of STAT1 at both 6 and 48 h (Figure 4B). Interestingly, at 6 h after challenge with *A. fumigatus* the phosphorylation of STAT1 in the *Mavs^-/-^* mice was not significantly different, although there was a slight but reproducible reduction (Figure 4B-C), which fits with our earlier observation that type I interferon mRNA levels were only mildly reduced at 3 h post-challenge with *A. fumigatus* (Figure 3A-B). In contrast, phosphorylation of STAT1 in the *Mavs^-/-^* mice was significantly decreased at 48 h post-challenge with *A. fumigatus* (Figure 4B-C), which corresponds with the significant decrease in type III interferon mRNA levels we observed at this time (Figure 3C). Taken together, these results strongly support the conclusion that MAVS signaling is critical for the late expression of type III interferons in response to *A. fumigatus* instillation and initiation of the interferon response that is necessary for host resistance against *A. fumigatus* infection.

**Figure 4.**
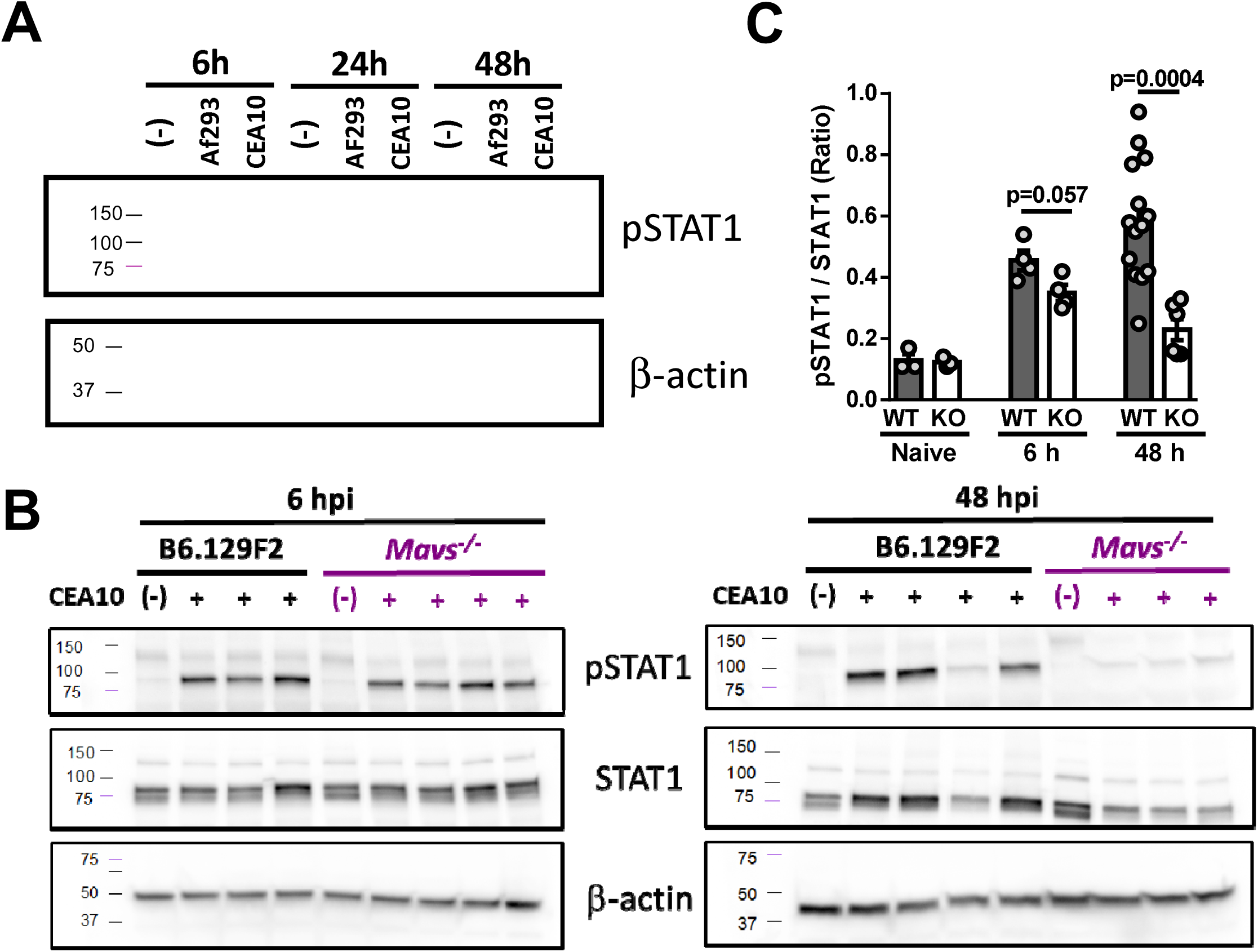
***Mavs* mice have decreased STAT1 phosphorylation after *Aspergillus fumigatus* challenge at later times. (A)** C57BL/6J mice were challenged with 4×10 conidia of CEA10 or Af293. Lungs were collected at 6, 24, and 48 hours post-inoculation and total protein was extracted from whole lungs. Protein expression and phosphorylation was determined by Western blot analysis. **(B)** B6.129F2 and *Mavs* mice were challenged with 4×10 conidia of CEA10. Lungs were collected at 6- or 48-hours post-inoculation and total protein was extracted from whole lungs. Protein expression and phosphorylation was determined by Western blot analysis. **(C)** Western blots from panel B were quantified by densitometry. Bars represent data means ± SEM with each dot representing individual mice. Data are representative of results from at least 2 independent experiments with at least 3 mice per group. Data were analyzed using a Mann-Whitney U-test.

### Alteration of the airway inflammatory milieu after *Aspergillus fumigatus* challenge in the absence of MAVS-dependent signaling

While our data demonstrates that all the classes of interferons are decreased to vary degrees after pulmonary *A. fumigatus* challenged, we next wanted to more broadly understand the cytokine and chemokine response in the airways of *Mavs^-/-^* mice. To achieve this, we challenged B6.129F2 and *Mavs^-/-^* mice with 4×10^7^ conidia of *A. fumigatus*. At 12 and 48 h after challenge with *A. fumigatus*, the BALF was analyzed using a 32-plex Milliplex cytokine assay. Numerous cytokines and chemokines--TNFα, IL-17A, CCL2, and CCL4--were expressed in a manner that was completely independent of MAVS signaling (Figure 5). Interestingly, IL-1α, IL-1β CXCL1, and CXCL2 secretion were significantly greater in the absence of MAVS signaling (Figure 5). In contrast, IL-12p70, CXCL9, and CXCL10 secretion was significantly impaired in the absence of MAVS signaling, particularly at 48 h post-challenge despite the large increase in fungal burden (Figure 5). These data suggest that MAVS-dependent signaling is critical for induction and/or maintenance of these cytokines.

**Figure 5.**
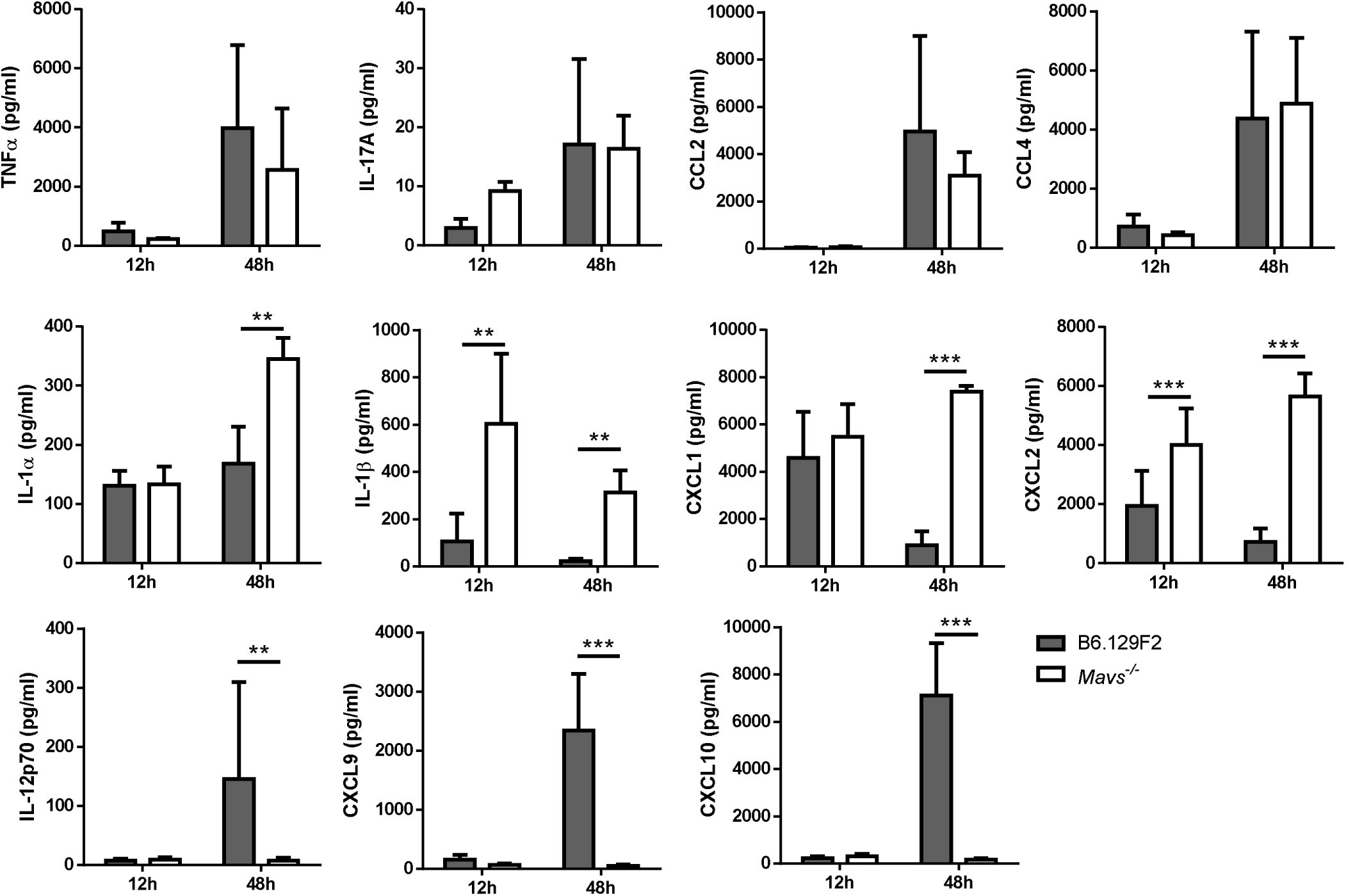
***Mavs* mice have an altered inflammatory milieu after *Aspergillus fumigatus* challenge in the airways.** B6.129F2 and *Mavs* mice were challenged with 4×10 conidia of CEA10. Bronchoalveolar lavage fluid (BALF) was collected 12 or 48h post-inoculation. Cytokine and chemokine levels were determined using a Milliplex Mouse Cytokine & Chemokine 32-plex (Millipore). Bars represent data means ± SEM. Data are representative of results 2 independent experiments with 4-6 mice per group. Data were analyzed using a Mann-Whitney U-test (* p < 0.05; ** p < 0.01; *** p < 0.001).

### *Cxcl9* and *Cxcl10* mRNA expression after pulmonary *Aspergillus fumigatus* challenge requires both type I and type III interferons

The marked decrease in CXCL9 and CXCL10 expression in the absence of MAVS following *A. fumigatus* challenge was quite striking, because both CXCL9 and CXCL10 are interferon-inducible chemokines (64). To test the role of type I and type III interferons in the induction of CXCL9 and CXCL10 expression, we challenged C57BL/6, *Ifnar^-/-^*, *Ifnlr^-/-^*, and *Ifnar^-/-^ Ifnlr^-/-^* (DKO) mice with 4×10^7^ conidia of *A. fumigatus*. Forty-eight hours later, lungs were collected for quantitative RT-PCR analysis to assess *Cxcl9* and *Cxcl10* mRNA expression patterns. Both *Ifnar-* and *Ifnlr-*deficient mice had a significant decrease in their expression of both *Cxcl9* (Figure 6A) and *Cxcl10* (Figure 6B). In general, the defect in *Cxcl9* and *Cxcl10* mRNA expression was greater in the absence of type III interferon signaling, particularly regarding *Cxcl9* expression. Importantly, the *Ifnar^-/-^ Ifnlr^-/-^* (DKO) mice had the most severe defect in *Cxcl9* and *Cxcl10* expression (Figure 6). Overall, these data demonstrate that type I and type III interferon signaling are essential for the expression of both *Cxcl9* and *Cxcl10*.

**Figure 6.**
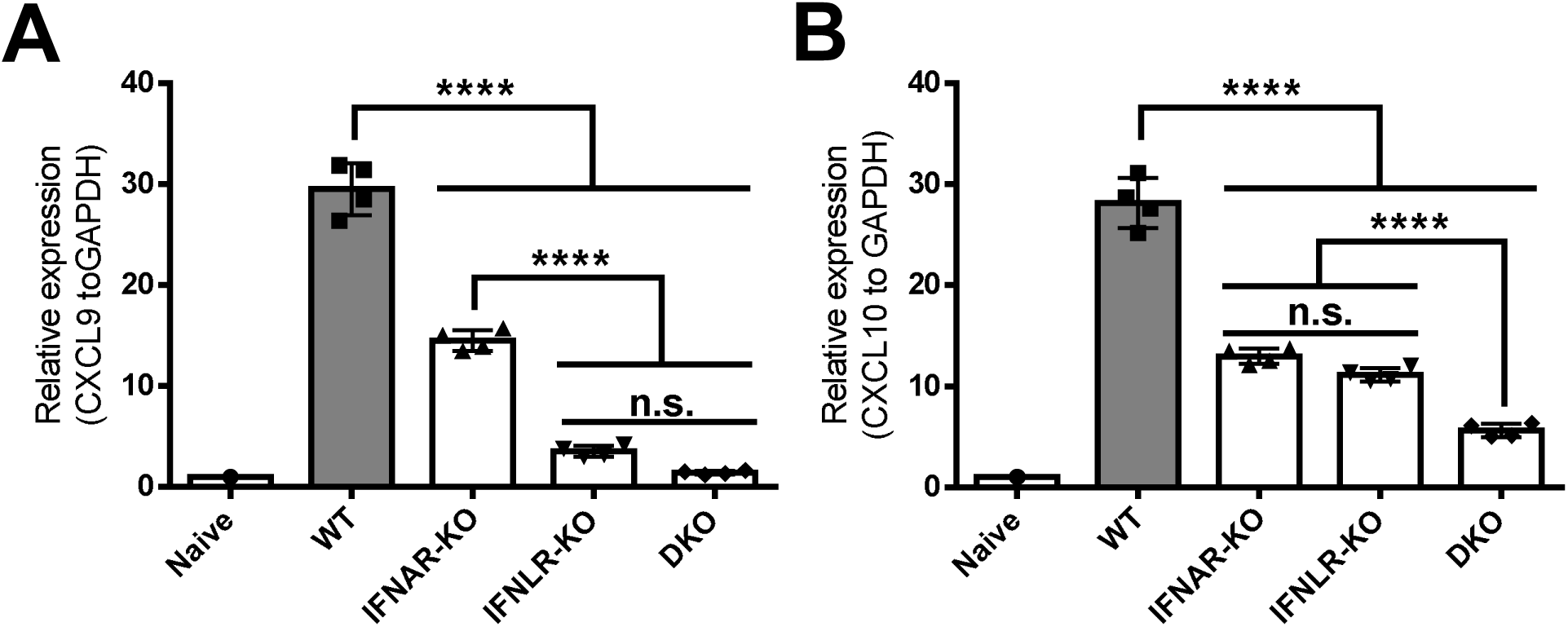
**Expression of *Cxcl9* and *Cxcl10* are dependent on IFN signaling.** C57BL/6J, *Ifnar*, *Ifnlr*, and *Ifnar Ifnlr* (DKO) mice were challenged with 4×10 conidia of CEA10. Lungs were collected 48h post-inoculation and total RNA was extracted. Gene expression as determined by quantitative reverse transcription polymerase chain reaction (qRT-PCR) using TaqMan probes for *Cxcl9* **(A)** and *Cxcl10*, which were normalized to *Gapdh* expression. Bars represent data means ± SEM with each symbol representing individual mice. Data are representative of results from at least 2 independent experiments with at least 4 mice per infected group. Data were analyzed using a Mann-Whitney U-test (n.s. = not significant; **** p < 0.0001).

### MDA5/MAVS-dependent signaling is required for host resistance against *Aspergillus fumigatus* infection

These previous data suggest that MDA5 can sense *A. fumigatus* RNA to drive an interferon response, which could be critical in vitality sensing following an *in vivo* challenge with *A. fumigatus*. Thus, we wanted to test the role of MDA5/MAVS signaling in host resistance against respiratory challenge with the human fungal pathogen, *A. fumigatus*. To globally assess the role of the RLR family in antifungal immunity, we initially challenged B6.129F2 mice and *Mavs^-/-^* mice with 4×10^7^ conidia of the CEA10 strain, since MAVS is the central signaling adaptor for both RIG-I and MDA5-mediated responses (65–68). We monitored the survival of the

*A. fumigatus* challenged mice over the next two weeks. *Mavs-*deficient mice were more susceptible to pulmonary challenge with *A. fumigatus* than wild-type, B6/129F2 mice (Figure 7A; Mantel-Cox log rank test, p = 0.0001). Additionally, susceptibility of the *Mavs^-/-^* mice to *A. fumigatus* challenge is not dependent on the strain of *A. fumigatus* used because *Mavs^-/-^* mice challenged with the Af293 strain were also highly susceptible to infection (Supplemental Figure 1; Mantel-Cox log rank test, p < 0.0001).

**Figure 7.**
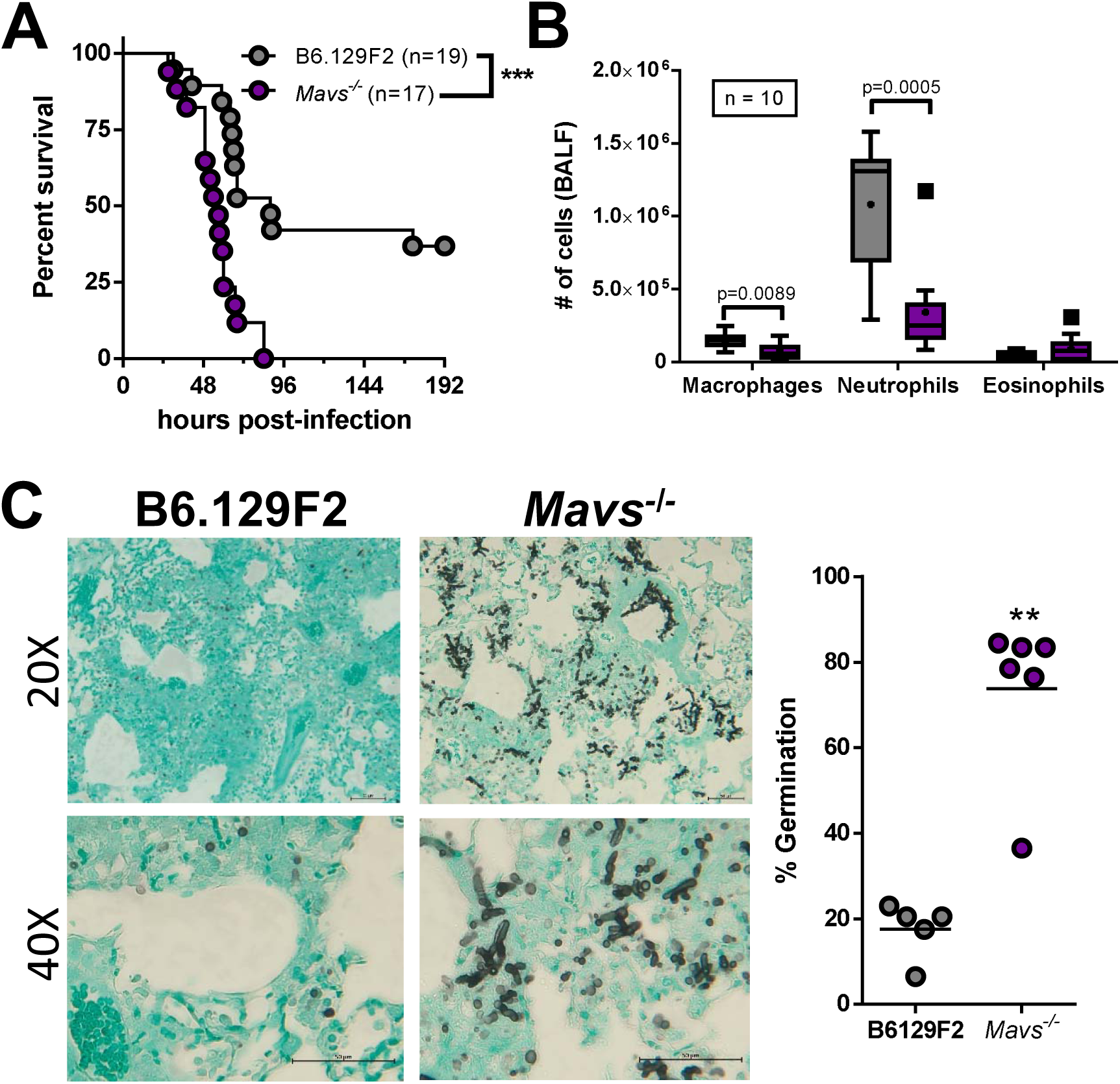
***Mavs* mice are highly susceptible to *Aspergillus fumigatus.*** B6.129F2 and *Mavs* mice were challenged with 4×10 conidia of CEA10. **(A)** Survival analysis in immune-competent wild-type and knock-out mice were tracked over the first days. Statistical significance was assessed using a Mantel-Cox log rank test (***p = 0.0001). **(B)** At 36 hpi, mice were euthanized and BALF collected for quantification of macrophage and neutrophil recruitment to the airways. Data are pooled from 2 independent experiments for a total of 10 mice per group. Data are presented as box-and-whisker plots with Tukey whiskers and outliers displayed as dots. Statistical significance was determined using a Mann-Whitney U test. **(C)** At 40 hpi mice were euthanized and lungs saved for histological analysis. Formalin-fixed lungs were paraffin embedded, sectioned, and stained with GMS for analysis by microscopy. Representative lung sections from are shown to the left using the 20x and 40x objective. *A. fumigatus* germination rates were determined by microscopically counting both the number of conidia and number of germlings in GMS-stained sections. Data from 2 independent experiments with 4-6 mice per group. Statistical significance was determined using a Mann-Whitney U test (**p < 0.01).

It is widely acknowledged that neutrophils and macrophages are critical antifungal effector cells for clearing *A. fumigatus* from the lungs. Thus, we assessed inflammatory cell accumulation in the airways via differential microscopic counting of cytospins stained with Diff-Quik from the BALF. The increased susceptibility of *Mavs-*deficient mice to *A. fumigatus* challenge was associated with decreased accumulation of macrophages and neutrophils in the airways compared with B6.129F2 mice at 40 h post-challenge (Figure 7B). Next, we assessed fungal growth and tissue invasion in the lung by histological analysis at 40 h after conidial instillation. Strikingly, GMS staining of lung tissue from *Mavs*-deficient mice revealed the presence of germinating *A. fumigatus* conidia at 40 h that was not observed to the same extent in B6.129F2 mice (Figure 7C). When the presence of germinating *A. fumigatus* conidia was quantified, wild-type B6.129F2 mice displayed low levels of fungal germination (17.6% ± 2.9) compared with *Mavs^-/-^* mice in which most of the fungal material had germinated (73.8% ± 7.6) by 40 h post-challenge (Figure 7C). B6.129F2 mice had very few areas of fungal material throughout their lungs, but it was associated with robust cellular infiltration and tissue destruction (Figure 7C).

To specifically address which RLR sensor is involved in regulating the innate anti-fungal immune response against *A. fumigatus* we chose to assess the role of MDA5 because of its role in the *in vitro* sensing of *A. fumigatus* RNA (Figure 2). Specifically, we challenged C57BL/6J and *Ifih1^-/-^* mice with 4×10^7^ conidia of *A. fumigatus*. We monitored the survival of the *A. fumigatus* challenged mice over the next ten days. Like the *Mavs*-deficient mice, *Ifih1-*deficient mice were more susceptible to pulmonary challenge with *A. fumigatus* than wild-type C57BL/6J mice (Figure 8A; Mantel-Cox log rank test, p = 0.0004). Analogous to what we observed in the *Mavs-*deficient mice, the increased susceptibility of *Ifih1-*deficient mice to *A. fumigatus* challenge was associated with decreased accumulation of neutrophils in the airways compared with C57BL/6J mice at 40 h post-challenge (Figure 8B). GMS staining of lung tissue from *Ifih1*- deficient mice revealed the presence of germinating *A. fumigatus* conidia at 42 h that was not observed to the same extent in C57BL/6J mice (Figure 8C) similar to our observation in the *Mavs-*deficient mice.

**Figure 8.**
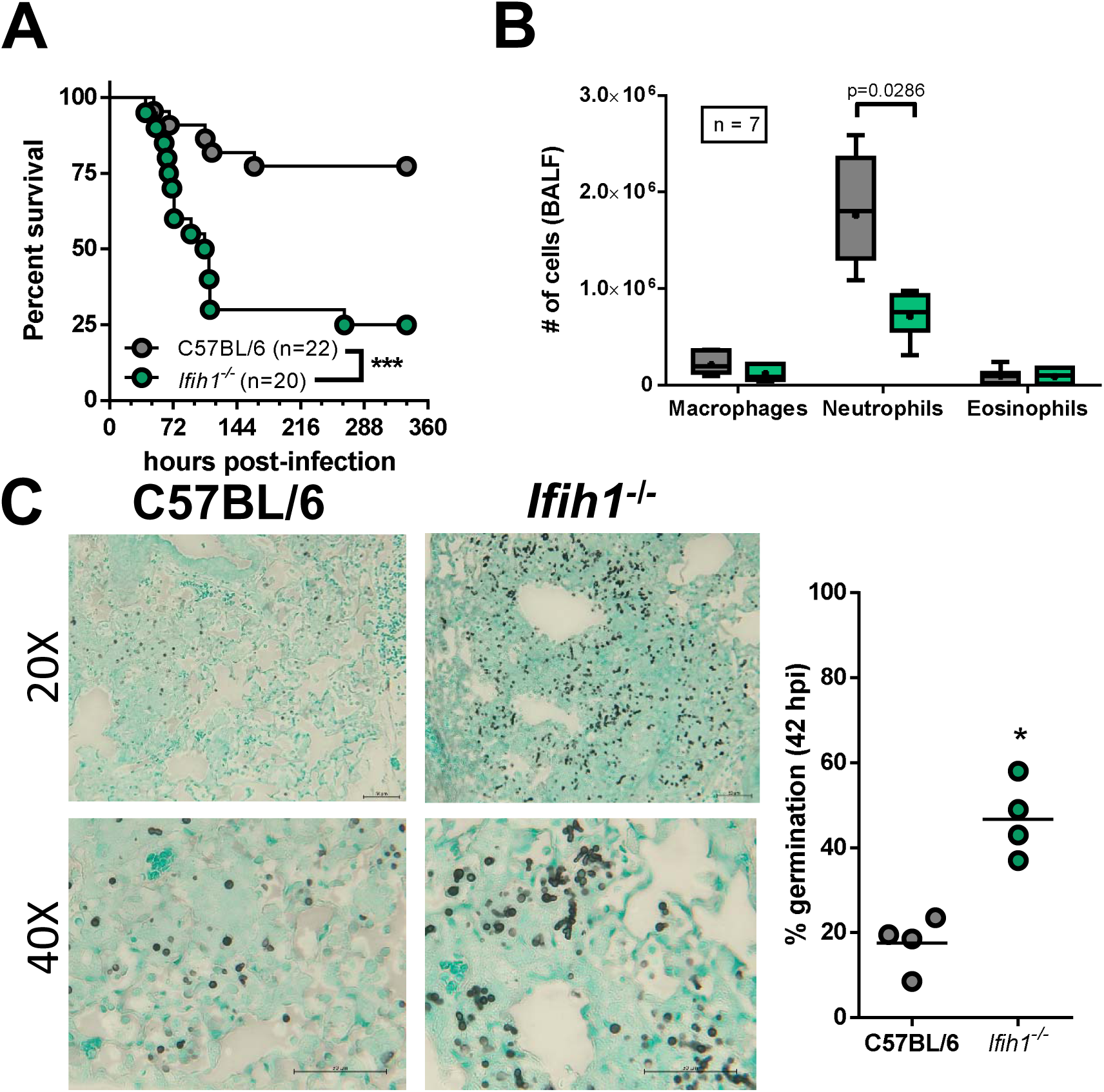
***Ifih1* mice are highly susceptible to *Aspergillus fumigatus.*** C57BL/6J and *Ifih1* mice were challenged with 4×10 conidia of CEA10. **(A)** Survival analysis in immune-competent wild-type and knock-out mice were tracked over the first 15 days. Statistical significance was assessed using a Mantel-Cox log rank test (***p = 0.0004). **(B & E)** At 36 hpi, mice were euthanized and BALF collected for quantification of macrophage and neutrophil recruitment to the airways. Data from 2 independent experiments for a total of 7 mice per group. Data are presented as box-and-whisker plots with Tukey whiskers and outliers displayed as dots. Statistical significance was determined using a Mann-Whitney U test. **(C & F)** At 42 hpi mice were euthanized and lungs saved for histological analysis. Formalin-fixed lungs were paraffin embedded, sectioned, and stained with GMS for analysis by microscopy. Representative lung sections from are shown to the right using the 20x and 40x objective. *A. fumigatus* germination rates were determined by microscopically counting both the number of conidia and number of germlings in GMS-stained sections. Data from 2 independent experiments with 4-6 mice per group. Statistical significance was determined using a Mann-Whitney U test (*p < 0.05; **p < 0.01).

While immune competent animal models have provided important insights into the host immune pathways necessary for antifungal control (11, 13, 56, 69–71), we wanted to assess the importance of MDA5/MAVS signaling in a clinically relevant immune compromised model of IPA. For this we utilized the triamcinolone model of IPA (72). Briefly, C57BL/6J and *Ifih1^-/-^* mice were immunosuppressed with triamcinolone and then subjected to challenge with 10^6^ conidia of the Af293 strain of *A. fumigatus*. At 72 h post-inoculation lungs were collected for histological analysis for fungal growth. Interestingly, *Ifih1^-/-^* mice had a significantly greater number of fungal lesions per lung section than C57BL/6J mice (Supplemental Figure 2A). Moreover, the fungal lesions appear to be larger and contain more fungi (Supplemental Figure 2A). Additionally, the BALF from *Ifih1^-/-^* challenged mice had higher levels of LDH activity (Supplemental Figure 2B) and albumin (Supplemental Figure 2C), which are indicators of lung damage and leakage, respectively, than C57BL/6J challenged mice. Taken together, these results strongly support the conclusion that MDA5/MAVS signaling is critical for host resistance against *A. fumigatus* in an immune competent murine model and critical for disease progression in a corticosteroid immune compromised murine model.

### Neutrophil antifungal effector functions are decreased in the absence of MDA5

To determine whether MDA5/MAVS signaling regulates antifungal effector cell functions, we used FLARE conidia challenge of C57BL/6 or *Ifih1^-/-^* mice to assess neutrophil-mediated conidial uptake and killing. FLARE conidia encode a fungal viability indicator (DsRed) and contain a tracer fluorophore (Alexa Fluor 633) (56). For this study we generated a novel FLARE strain in the CEA10 strain background (Supplemental Figure 3). FLARE conidia emit two fluorescence signals (DsRed and Alexa Fluor 633) when the conidia are alive, but only emit a single fluorescence signal (Alexa Fluor 633) when the conidia are dead. This approach allows us to determine the frequency of conidia-engaged immune cells that contain either live or dead fungal cells in the BAL and lungs. The frequency of conidia-engaged neutrophils was similar between the C57BL/6 and *Ifih1^-/-^* mice (Figure 9A). However, the frequency of conidia-engaged neutrophils that contain live conidia was increased among *Ifhi1^−/−^* mice compared with C57BL/6J mice, indicating a defect in conidial killing (Figure 8B). Specifically, the frequency of neutrophils that contain live conidia was 1.9-fold and 2.1-fold higher for BAL and lung fluid of *Ifih1^−/−^* mice compared with their C57BL/6J counterparts (Figure 9B). Thus, these data indicate that neutrophil-mediated antifungal killing of *A. fumigatus* conidia requires MDA5/MAVS activation and signaling.

**Figure 9.**
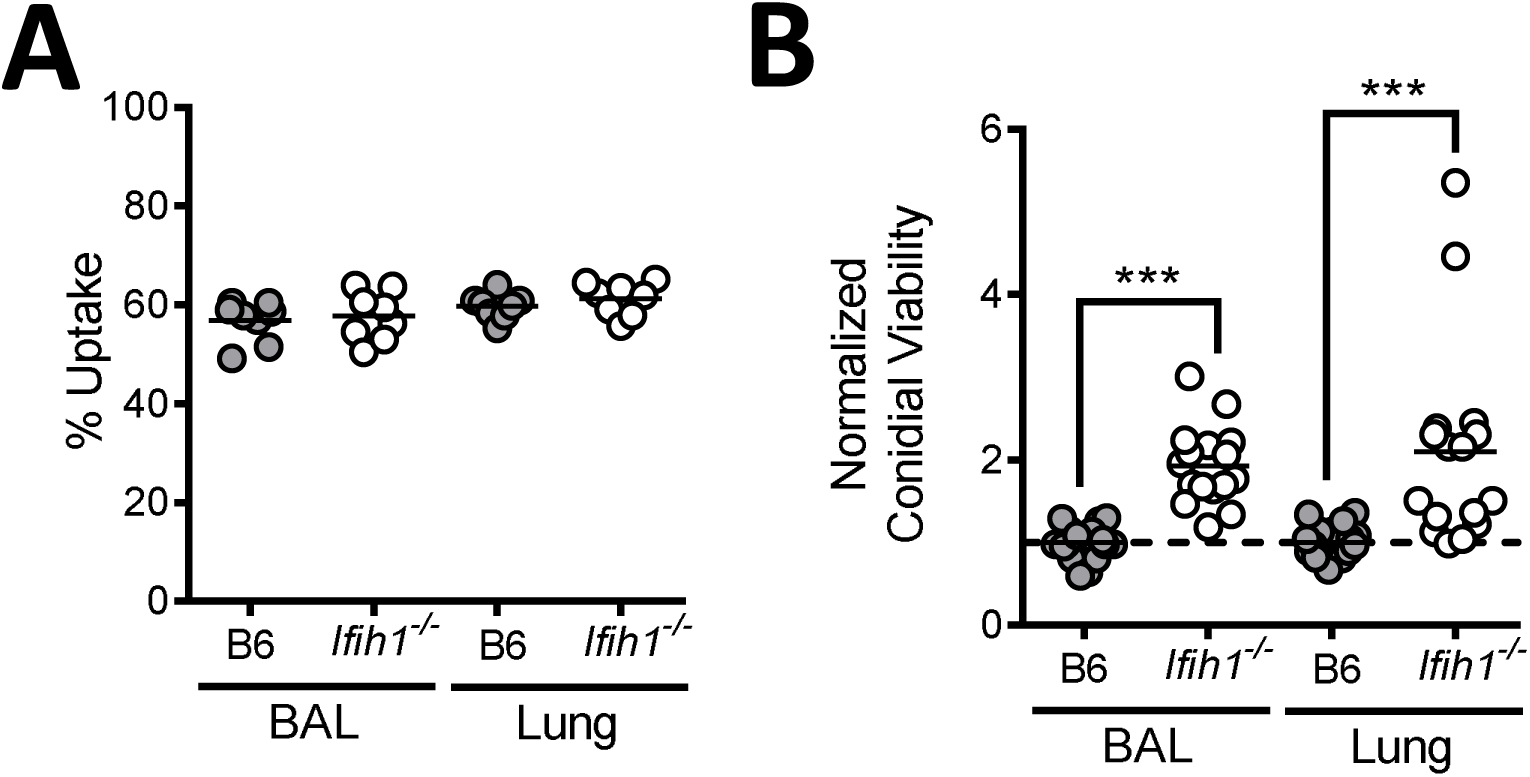
***Ifih1^-/-^* mice have decrease antifungal killing by neutrophils *in vivo.*** C57BL/6J and *Ifih1^-/-^* mice were challenged with 3x10^7^ conidia of CEA10 fluorescent *Aspergillus* reporter (FLARE). Bronchoalveolar lavage (BAL) fluid and lungs were harvested at 48 hours after infection and fungal uptake and viability were analyzed with flow cytometry. Fungal uptake **(A)** and viability **(B)** in neutrophils were measured. Data pooled from 2 independent experiments with 7-9 mice per group. Statistical significance was determined using a Mann-Whitney U test (***p < 0.01).

## DISCUSSION

It is now well appreciated that the host inflammatory response induced by *A. fumigatus* is tightly tuned to the virulence of the individual strain under study (26–30). One key determinant for sensing the threat posed by filamentous fungi is changes in fungal cell wall composition. Specifically, detection of β-1,3-glucan linked polysaccharides by Dectin-1 (*Clec7a*) occurs only upon conidial swelling and germling formation, which are the earliest steps of fungal growth (53, 59). Interestingly, our data demonstrate that while heat-killed swollen conidia of *A. fumigatus* can induce the secretion of pro-inflammatory cytokines, such as TNFα, they do not drive the secretion of IFNα and CXCL10 (Figure 1). These data suggest that heat sensitive fungal factors other than cell wall polysaccharides are responsible for driving the interferon response following respiratory *A. fumigatus* challenge. Consistent with this observation, individual polysaccharides commonly found in the *A. fumigatus* cell wall are not sufficient to induce an interferon response (73). Thus, other less studied fungal traits, that appear to require live fungi, are needed to induce a protective interferon response.

Intriguingly, during bacterial infections one key virulence trait sensed by the host immune system to tune the inflammatory response is the ability of the bacteria to resist killing and remain viable. Two crucial inflammatory pathways that signal bacterial vitality to the host are inflammasome secretion of IL-1β and an interferon response through TRIF and cGAS/STING signaling (33, 34, 74). Following respiratory challenge with *A. fumigatus* both IL-1β and type I/III interferons have been shown to be critical for antifungal immunity (35, 39). While we and others have previously demonstrated that inflammasomes are essential for the IL-1β response after respiratory challenge with *A. fumigatus* (35–38), the innate pattern-recognition receptor(s) leading to type I/III interferon expression have not been elucidated. Moreover, fungal molecules that signal vitality to the host are not yet defined.

In bacteria one critical PAMP associated with vitality (vita-PAMP) is RNA. Using an *E. coli* infection model, bacterial RNA was shown to be sufficient to drive both inflammasome-dependent IL-1β secretion and TRIF-dependent IFNβ secretion (33). Heat-killing of *E. coli* drove the loss of RNA and its subsequent inflammatory response (33). Several innate RNA sensors have been identified in mammalian hosts, including TLR3 (75), TLR7 (76), MDA5 (77), and RIG-I (78). TLR3 and TLR7 signal through TRIF (79), while MDA5 and RIG-I signal through MAVS (65–68) to drive the interferon response. In our study, RNase III treatment of total RNA isolated from *A. fumigatus* demonstrates that dsRNA from *A. fumigatus* is sufficient to drive an interferon response (Figure 2B). TLR3 and MDA5 are both receptors for dsRNA (60, 75, 78). Importantly, mice lacking *Mavs* displayed a markedly reduced type III interferon (IL-28/IFNλ), CXCL9, and CXCL10 secretion, but only a moderate defect in type I interferons (Figure 4). In contrast, the expression of *Ifna1* and *Ifnb1* are regulated by both TRIF (46) and TLR3 (45). Interestingly, the role of TRIF appears to be restricted to the non-hematopoietic compartment (46), likely epithelial cells (80). Taken together these data suggest the type I interferon response following live *A. fumigatus* challenge is primarily driven by TLR3/TRIF-dependent signaling, while the type III interferon response is primarily drive by MDA5/MAVS-dependent signaling.

Our data raises an interesting conundrum of how RNA from live *A. fumigatus* enters host cell cytoplasm to drive MDA5/MAVS activation. One potential mechanism for the translocation of RNA from the phagosome to the cytosol is damage to the phagosomal membrane and the leakage of phagosomal contents to cytosol. In their original work describing vita-PAMPs Blander and colleagues found that phagosomes containing *E. coli* exhibited intrinsic leakiness, which enabled RNA from the live *E. coli* to enter the cytosol and activate cytosolic PRRs (33). However, phagosome leakiness is not unique to these bacterial systems as this has been previously described for particles such as beads and crystals that induce phagosome destabilization (81, 82). Interestingly, fungal pathogens can also drive phagosome destabilization and rupture leading to the activation of inflammasomes and pyroptosis (26, 83). In contrast to the passive nature of phagosomal leakage, SIDT2 has been shown to actively transport of dsRNA from the endosome to the cytosol for detection by RLRs, which was necessary for the IFN response to Poly(I:C), ECMV, and HSV-1 (84). Another alternative pathway may be extracellular vesicles. Fungal extracellular vesicles contain a variety of cargos including RNA, polysaccharides, and enzymes (85). Since extracellular vesicles possess the lipid bilayer structure like liposomes, they could deliver their contents to the cytosol via membrane fusion. Interestingly, extracellular vesicles from *Mycobacterium tuberculosis* have been shown to be sufficient for the activation of RIG-I (86, 87). Studies examining how *A. fumigatus* RNA found in the phagosomal compartment gains access to the cytoplasm to mediate the activation of cytosolic PRRs to drive the enhanced vita-PAMP response are on-going.

The purpose of vitality sensing is to drive robust inflammation that is in line with the threat posed by the invading pathogen. Neutrophils are well known to be critical antifungal effector cells needed for the prevention of fungal germination and clearance of *A. fumigatus* from the lungs (20). Our results demonstrate that MDA5/MAVS-dependent signaling is necessary for both the optimal accumulation (Figures 7 and 8) and activation of neutrophils in *Aspergillus* challenged murine airways (Figure 9). Following pulmonary challenge with *A. fumigatus* it has been established that both type I and type III interferons are essential for host resistance through enhancing neutrophil ROS production and antifungal effector functions (39). Our FLARE data support these previous observations as neutrophils in both the lungs and airways of *Ifih1-* deficient mice are significantly impaired in their antifungal killing capacity at 48 h after challenge (Figure 9), which corresponds with a time when type III interferons are nearly absent in mice lacking MDA5/MAVS signaling. Decreased accumulation of neutrophils in the *Mavs-* and *Ifih1*-deficient mice did not correlate with classical neutrophil chemoattractants, such as CXCR2 ligands, that are required to maintain host resistance against *A. fumigatus* (36, 70, 88). Rather, in the absence of *Mavs* and *Ifih1* there was a marked decrease in CXCL9 and CXCL10, which are both ligands for CXCR3. Recently, CXCR3 has been suggested to be a potential chemoattractant for neutrophils in certain systems (89, 90). However, *Cxcr3-*deficient mice appear to recruit normal numbers of neutrophils following *A. fumigatus* challenge (25). Thus, why *Mavs-* and *Ifih1*-deficient mice accumulate few neutrophils in their lungs following challenge warrants further research.

Type I interferon response can have both beneficial and detrimental effects to the host. For example, during LCMV Armstrong infection type I interferons drive host resistance (91), but excessive type I interferon levels found following LCMV Clone 13 infection promote virus persistence and immunopathology (92–94). Thus, the optimal tuning of the magnitude of the type I interferon response can dictate disease outcome following viral infection. The role of type I interferons during fungal infections has now been studied in several model systems, demonstrating a similar dichotomy. Following *A. fumigatus* challenge both type I and type III interferons are essential for host resistance against invasive aspergillosis through enhancing neutrophil ROS production and antifungal effector functions (24, 39). Moreover, augmentation of the type I interferon in X-CGD mice with invasive aspergillosis following treatment with Poly(I:C) improved disease outcomes (95). Similarly, during *Cryptococcus neoformans* treatment with Poly(I:C) drove type I interferon expression and iron limitation which corresponded with improved disease outcomes (96, 97). However, opposing roles of type I interferon signaling in T cell polarization following *C. neoformans* challenge have been observed, either promoting protective Th1 cells (98) or non-protective Th2 cells (99). After *Histoplasma capsulatum* challenge type I interferon signaling is essential for host resistance (100). The role of type I interferons during *Candida* spp. infections appear to be more complex. During *Candida glabrata* infection type I interferons promote fungal persistence through dampening nutritional immunity (101–103). Following *C. albicans* challenge, type I interferons either promote host resistance (49, 50, 104) or promote immunopathological immune responses which enhance disease (105). Thus, much remains to be learned about how interferons regulate infection outcomes during fungal infections. In this regard, how the detection of cell wall PAMPs, like β-1,3-glucans, by Dectin-1 (47, 50, 51) and fungal RNA, through TLRs (47, 48, 100, 103) and RLRs (63), coordinate the type I and type III interferon responses needed to be further explored. vita-PAMPs appear to be essential for the optimal induction the RNA driven interferon responses.

While the role of RLRs in host immunity was originally associated with antiviral innate immune responses (reviewed by (42)), recent studies have begun to expand the role of RLRs in immunity to other pathogens, including *C. albicans, Listeria monocytogenes*, *Mycobacterium tuberculosis*, and *Plasmodium falciparum* (40, 63, 86, 106–109). Our study adds *A. fumigatus* to the list of non-viral pathogens which can activate the RLR family. Further studies aiming to decipher the cellular and molecular mechanisms by which *A. fumigatus* enhances antifungal immunity through MDA5 activation are expected to inform us how the host uses these vita-PAMPs to sense the threat posed by fungal pathogens. Understanding how best to alter the host to the treat a pathogen poses will enable us to develop strategies to harness the inflammatory response for prophylactic or therapeutic gain in vulnerable populations. Interestingly, exogenous Poly(I:C) treatment prophylactically improves outcomes of invasive aspergillosis in *gp91^phox^*-deficient mice, particularly when challenged with resting conidia rather than germlings (95). Our previous work has highlighted that germlings induced a highly inflammatory response dependent on IL-1α (26) and LTB_4_ (27); thus, understanding the interplay and activation of the type I/III on IL-1α (26) and LTB_4_ inflammatory response necessary to control both the conidial and hyphal forms will essential for completely understanding host resistance and therapeutic options against *A. fumigatus*. Finally, our study raises the possibility that MDA5/MAVS and type I/III interferon signaling in critical for preventing invasive aspergillosis in vulnerable human patient populations.

## AUTHOR CONTRIBUTIONS

Conceived and designed the experiments: XW, AKC, AR, RC, JJO. Performed the experiments: XW, AKC, VE, KL, WC, SD. Analyzed the data: XW, AKC, VE, KL, WC, AR, JJO. Wrote the paper: XW, JJO.

XW and AKC are co-first authors: AKC initiated these studies before graduating, while XW picked up and completed these studies and helped write the manuscript, which warranted listing XW first.

## ACKNOWLEDGEMENTS

Thank you to Dr. Brent Berwin (Geisel School of Medicine at Dartmouth), Dr. David Leib (Geisel School of Medicine at Dartmouth), and Dr. Tobias Hohl (Memorial Sloan Kettering Cancer Center) for helpful discussion on this project and manuscript. Thank you to Stacy Ceron for assistance in generation of the primary murine fibroblast cultures.

## Notes

**Conflict of Interests:** The authors have declared that no conflict of interest exists.

1 Research in this study was supported in part by institutional startup funds to JJO in part through the Dartmouth Lung Biology Center for Molecular, Cellular, and Translational Research grant P30 GM106394 (PI: Bruce A. Stanton) and Center for Molecular, Cellular and Translational Immunological Research grant P30 GM103415 (PI: William R. Green). JJO was partially supported by a Munck-Pfefferkorn Award from Dartmouth College and NIH R01 AI139133 grant. Work by AR was partially funded by NIH R01 AI114647 grant. AR and RAC are Investigators in the Pathogenesis of Infectious Diseases supported by the Burroughs Wellcome Fund. The funders had no role in the preparation or publication of the manuscript.

### Competing Interest Statement

The authors have declared no competing interest.

## REFERENCES

1. Chazalet, V., J.P. Debeaupuis, J. Sarfati, J. Lortholary, P. Ribaud, P. Shah, M. Cornet, H. Vu Thien, E. Gluckman, G. Brucker, and J.P. Latge. 1998. Molecular typing of environmental and patient isolates of Aspergillus fumigatus from various hospital settings. J Clin Microbiol 36: 1494–1500.

2. Goodley, J.M., Y.M. Clayton, and R.J. Hay. 1994. Environmental sampling for aspergilli during building construction on a hospital site. J Hosp Infect 26: 27–35.

3. Hospenthal, D.R., K.J. Kwon-Chung, and J.E. Bennett. 1998. Concentrations of airborne Aspergillus compared to the incidence of invasive aspergillosis: lack of correlation. Med Mycol 36: 165–168.

4. Brown, G.D., D.W. Denning, N.A. Gow, S.M. Levitz, M.G. Netea, and T.C. White. 2012. Hidden killers: human fungal infections. Sci Transl Med 4: 165rv113.

5. Baddley, J.W., D.R. Andes, K.A. Marr, D.P. Kontoyiannis, B.D. Alexander, C.A. Kauffman, R.A. Oster, E.J. Anaissie, T.J. Walsh, M.G. Schuster, J.R. Wingard, T.F. Patterson, J.I. Ito, O.D. Williams, T. Chiller, and P.G. Pappas. 2010. Factors associated with mortality in transplant patients with invasive aspergillosis. Clin Infect Dis 50: 1559–1567.

6. Steinbach, W.J., K.A. Marr, E.J. Anaissie, N. Azie, S.P. Quan, H.U. Meier-Kriesche, S. Apewokin, and D.L. Horn. 2012. Clinical epidemiology of 960 patients with invasive aspergillosis from the PATH Alliance registry. J Infect 65: 453–464.

7. Garcia-Vidal, C., A. Upton, K.A. Kirby, and K.A. Marr. 2008. Epidemiology of invasive mold infections in allogeneic stem cell transplant recipients: biological risk factors for infection according to time after transplantation. Clin Infect Dis 47: 1041–1050.

8. Upton, A., K.A. Kirby, P. Carpenter, M. Boeckh, and K.A. Marr. 2007. Invasive aspergillosis following hematopoietic cell transplantation: outcomes and prognostic factors associated with mortality. Clin Infect Dis 44: 531–540.

9. Thompson, G.R., 3rd and T.F. Patterson. 2008. Pulmonary aspergillosis. Semin Respir Crit Care Med 29: 103–110.

10. Bongomin, F., S. Gago, R.O. Oladele, and D.W. Denning. 2017. Global and Multi-National Prevalence of Fungal Diseases-Estimate Precision. J Fungi (Basel) 3: 67.

11. Rieber, N., R.P. Gazendam, A.F. Freeman, A.P. Hsu, A.L. Collar, J.A. Sugui, R.A. Drummond, C. Rongkavilit, K. Hoffman, C. Henderson, L. Clark, M. Mezger, M. Swamydas, M. Engeholm, R. Schule, B. Neumayer, F. Ebel, C.M. Mikelis, S. Pittaluga, V.K. Prasad, A. Singh, J.D. Milner, K.W. Williams, J.K. Lim, K.J. Kwon-Chung, S.M. Holland, D. Hartl, T.W. Kuijpers, and M.S. Lionakis. 2016. Extrapulmonary Aspergillus infection in patients with CARD9 deficiency. JCI Insight 1: e89890.

12. Fisher, C.E., T.M. Hohl, W. Fan, B.E. Storer, D.M. Levine, L.P. Zhao, P.J. Martin, E.H. Warren, M. Boeckh, and J.A. Hansen. 2017. Validation of single nucleotide polymorphisms in invasive aspergillosis following hematopoietic cell transplantation. Blood 129: 2693–2701.

13. Cunha, C., F. Aversa, J.F. Lacerda, A. Busca, O. Kurzai, M. Grube, J. Loffler, J.A. Maertens, A.S. Bell, A. Inforzato, E. Barbati, B. Almeida, P. Santos e Sousa, A. Barbui, L. Potenza, M. Caira, F. Rodrigues, G. Salvatori, L. Pagano, M. Luppi, A. Mantovani, A. Velardi, L. Romani, and A. Carvalho. 2014. Genetic PTX3 deficiency and aspergillosis in stem-cell transplantation. N Engl J Med 370: 421–432.

14. Morgenstern, D.E., M.A. Gifford, L.L. Li, C.M. Doerschuk, and M.C. Dinauer. 1997. Absence of respiratory burst in X-linked chronic granulomatous disease mice leads to abnormalities in both host defense and inflammatory response to Aspergillus fumigatus. J Exp Med 185: 207–218.

15. Pollock, J.D., D.A. Williams, M.A. Gifford, L.L. Li, X. Du, J. Fisherman, S.H. Orkin, C.M. Doerschuk, and M.C. Dinauer. 1995. Mouse model of X-linked chronic granulomatous disease, an inherited defect in phagocyte superoxide production. Nat Genet 9: 202–209.

16. Grimm, M.J., R.R. Vethanayagam, N.G. Almyroudis, C.G. Dennis, A.N. Khan, A.C. D’Auria, K.L. Singel, B.A. Davidson, P.R. Knight, T.S. Blackwell, T.M. Hohl, M.K. Mansour, J.M. Vyas, M. Rohm, C.F. Urban, T. Kelkka, R. Holmdahl, and B.H. Segal. 2013. Monocyte- and macrophage-targeted NADPH oxidase mediates antifungal host defense and regulation of acute inflammation in mice. J Immunol 190: 4175–4184.

17. Bhatia, S., M. Fei, M. Yarlagadda, Z. Qi, S. Akira, S. Saijo, Y. Iwakura, N. van Rooijen, G.A. Gibson, C.M. St Croix, A. Ray, and P. Ray. 2011. Rapid host defense against Aspergillus fumigatus involves alveolar macrophages with a predominance of alternatively activated phenotype. PLoS One 6: e15943.

18. Espinosa, V., A. Jhingran, O. Dutta, S. Kasahara, R. Donnelly, P. Du, J. Rosenfeld, I. Leiner, C.C. Chen, Y. Ron, T.M. Hohl, and A. Rivera. 2014. Inflammatory monocytes orchestrate innate antifungal immunity in the lung. PLoS Pathog 10: e1003940.

19. Hohl, T.M., A. Rivera, L. Lipuma, A. Gallegos, C. Shi, M. Mack, and E.G. Pamer. 2009. Inflammatory monocytes facilitate adaptive CD4 T cell responses during respiratory fungal infection. Cell Host Microbe 6: 470–481.

20. Mircescu, M.M., L. Lipuma, N. van Rooijen, E.G. Pamer, and T.M. Hohl. 2009. Essential role for neutrophils but not alveolar macrophages at early time points following Aspergillus fumigatus infection. J Infect Dis 200: 647–656.

21. Bonnett, C.R., E.J. Cornish, A.G. Harmsen, and J.B. Burritt. 2006. Early neutrophil recruitment and aggregation in the murine lung inhibit germination of Aspergillus fumigatus Conidia. Infect Immun 74: 6528–6539.

22. Cohen, N.R., R.V. Tatituri, A. Rivera, G.F. Watts, E.Y. Kim, A. Chiba, B.B. Fuchs, E. Mylonakis, G.S. Besra, S.M. Levitz, M. Brigl, and M.B. Brenner. 2011. Innate recognition of cell wall beta-glucans drives invariant natural killer T cell responses against fungi. Cell Host Microbe 10: 437–450.

23. Loures, F.V., M. Rohm, C.K. Lee, E. Santos, J.P. Wang, C.A. Specht, V.L. Calich, C.F. Urban, and S.M. Levitz. 2015. Recognition of Aspergillus fumigatus hyphae by human plasmacytoid dendritic cells is mediated by dectin-2 and results in formation of extracellular traps. PLoS Pathog 11: e1004643.

24. Ramirez-Ortiz, Z.G., C.K. Lee, J.P. Wang, L. Boon, C.A. Specht, and S.M. Levitz. 2011. A nonredundant role for plasmacytoid dendritic cells in host defense against the human fungal pathogen Aspergillus fumigatus. Cell Host Microbe 9: 415–424.

25. Guo, Y., S. Kasahara, A. Jhingran, N.L. Tosini, B. Zhai, M.A. Aufiero, K.A.M. Mills, M. Gjonbalaj, V. Espinosa, A. Rivera, A.D. Luster, and T.M. Hohl. 2020. During Aspergillus Infection, Monocyte-Derived DCs, Neutrophils, and Plasmacytoid DCs Enhance Innate Immune Defense through CXCR3-Dependent Crosstalk. Cell Host Microbe doi: 10.1016/j.chom.2020.1005.1002.

26. Caffrey-Carr, A.K., C.H. Kowalski, S.R. Beattie, N.A. Blaseg, C.R. Upshaw, A. Thammahong, H.E. Lust, Y.W. Tang, T.M. Hohl, R.A. Cramer, and J.J. Obar. 2017. IL-1alpha is Critical for Resistance Against Highly Virulent Aspergillus fumigatus Isolates. Infect Immun 85: e00661–00617.

27. Caffrey-Carr, A.K., K.M. Hilmer, C.H. Kowalski, K.M. Shepardson, R.M. Temple, R.A. Cramer, and J.J. Obar. 2018. Host-Derived Leukotriene B4 Is Critical for Resistance against Invasive Pulmonary Aspergillosis. Front Immunol 8: 1984.

28. Caffrey, A.K. and J.J. Obar. 2016. Alarmin(g) the innate immune system to invasive fungal infections. Curr Opin Microbiol 32: 135–143.

29. Rizzetto, L., G. Giovannini, M. Bromley, P. Bowyer, L. Romani, and D. Cavalieri. 2013. Strain dependent variation of immune responses to A. fumigatus: definition of pathogenic species. PLoS One 8: e56651.

30. Rosowski, E.E., N. Raffa, B.P. Knox, N. Golenberg, N.P. Keller, and A. Huttenlocher. 2018. Macrophages inhibit Aspergillus fumigatus germination and neutrophil-mediated fungal killing. PLoS Pathog 14: e1007229.

31. Vance, R.E., R.R. Isberg, and D.A. Portnoy. 2009. Patterns of pathogenesis: discrimination of pathogenic and nonpathogenic microbes by the innate immune system. Cell Host Microbe 6: 10–21.

32. Blander, J.M. and L.E. Sander. 2012. Beyond pattern recognition: five immune checkpoints for scaling the microbial threat. Nat Rev Immunol 12: 215–225.

33. Sander, L.E., M.J. Davis, M.V. Boekschoten, D. Amsen, C.C. Dascher, B. Ryffel, J.A. Swanson, M. Muller, and J.M. Blander. 2011. Detection of prokaryotic mRNA signifies microbial viability and promotes immunity. Nature 474: 385–389.

34. Barbet, G., L.E. Sander, M. Geswell, I. Leonardi, A. Cerutti, I. Iliev, and J.M. Blander. 2018. Sensing Microbial Viability through Bacterial RNA Augments T Follicular Helper Cell and Antibody Responses. Immunity 48: 584–598.

35. Karki, R., S.M. Man, R.K. Malireddi, P. Gurung, P. Vogel, M. Lamkanfi, and T.D. Kanneganti. 2015. Concerted activation of the AIM2 and NLRP3 inflammasomes orchestrates host protection against Aspergillus infection. Cell Host Microbe 17: 357–368.

36. Caffrey, A.K., M.M. Lehmann, J.M. Zickovich, V. Espinosa, K.M. Shepardson, C.P. Watschke, K.M. Hilmer, A. Thammahong, B.M. Barker, A. Rivera, R.A. Cramer, and J.J. Obar. 2015. IL-1alpha signaling is critical for leukocyte recruitment after pulmonary Aspergillus fumigatus challenge. PLoS Pathog 11: e1004625.

37. Said-Sadier, N., E. Padilla, G. Langsley, and D.M. Ojcius. 2010. Aspergillus fumigatus stimulates the NLRP3 inflammasome through a pathway requiring ROS production and the Syk tyrosine kinase. PLoS One 5: e10008.

38. Moretti, S., S. Bozza, V. Oikonomou, G. Renga, A. Casagrande, R.G. Iannitti, M. Puccetti, C. Garlanda, S. Kim, S. Li, F.L. van de Veerdonk, C.A. Dinarello, and L. Romani. 2014. IL-37 inhibits inflammasome activation and disease severity in murine aspergillosis. PLoS Pathog 10: e1004462.

39. Espinosa, V., O. Dutta, C. McElrath, P. Du, Y.J. Chang, B. Cicciarelli, A. Pitler, I. Whitehead, J.J. Obar, J.E. Durbin, S.V. Kotenko, and A. Rivera. 2017. Type III interferon is a critical regulator of innate antifungal immunity Science Immunology 2: eaan5357.

40. Wu, J., L. Tian, X. Yu, S. Pattaradilokrat, J. Li, M. Wang, W. Yu, Y. Qi, A.E. Zeituni, S.C. Nair, S.P. Crampton, M.S. Orandle, S.M. Bolland, C.F. Qi, C.A. Long, T.G. Myers, J.E. Coligan, R. Wang, and X.Z. Su. 2014. Strain-specific innate immune signaling pathways determine malaria parasitemia dynamics and host mortality. Proc Natl Acad Sci U S A 111: E511–520.

41. Lazear, H.M., J.W. Schoggins, and M.S. Diamond. 2019. Shared and Distinct Functions of Type I and Type III Interferons. Immunity 50: 907–923.

42. Dixit, E. and J.C. Kagan. 2013. Intracellular pathogen detection by RIG-I-like receptors. Adv Immunol 117: 99–125.

43. Herbst, S., A. Shah, M. Mazon Moya, V. Marzola, B. Jensen, A. Reed, M.A. Birrell, S. Saijo, S. Mostowy, S. Shaunak, and D. Armstrong-James. 2015. Phagocytosis-dependent activation of a TLR9-BTK-calcineurin-NFAT pathway co-ordinates innate immunity to Aspergillus fumigatus. EMBO Mol Med 7: 240–258.

44. Ramaprakash, H., T. Ito, T.J. Standiford, S.L. Kunkel, and C.M. Hogaboam. 2009. Toll-like receptor 9 modulates immune responses to Aspergillus fumigatus conidia in immunodeficient and allergic mice. Infect Immun 77: 108–119.

45. Carvalho, A., A. De Luca, S. Bozza, C. Cunha, C. D’Angelo, S. Moretti, K. Perruccio, R.G. Iannitti, F. Fallarino, A. Pierini, J.P. Latge, A. Velardi, F. Aversa, and L. Romani. 2012. TLR3 essentially promotes protective class I-restricted memory CD8(+) T-cell responses to Aspergillus fumigatus in hematopoietic transplanted patients. Blood 119: 967–977.

46. de Luca, A., S. Bozza, T. Zelante, S. Zagarella, C. D’Angelo, K. Perruccio, C. Vacca, A. Carvalho, C. Cunha, F. Aversa, and L. Romani. 2010. Non-hematopoietic cells contribute to protective tolerance to Aspergillus fumigatus via a TRIF pathway converging on IDO. Cell Mol Immunol 7: 459–470.

47. Khan, N.S., P.V. Kasperkovitz, A.K. Timmons, M.K. Mansour, J.M. Tam, M.W. Seward, J.L. Reedy, S. Puranam, M. Feliu, and J.M. Vyas. 2016. Dectin-1 Controls TLR9 Trafficking to Phagosomes Containing beta-1,3 Glucan. J Immunol 196: 2249–2261.

48. Biondo, C., A. Malara, A. Costa, G. Signorino, F. Cardile, A. Midiri, R. Galbo, S. Papasergi, M. Domina, M. Pugliese, G. Teti, G. Mancuso, and C. Beninati. 2012. Recognition of fungal RNA by TLR7 has a nonredundant role in host defense against experimental candidiasis. Eur J Immunol 42: 2632–2643.

49. Biondo, C., G. Signorino, A. Costa, A. Midiri, E. Gerace, R. Galbo, A. Bellantoni, A. Malara, C. Beninati, G. Teti, and G. Mancuso. 2011. Recognition of yeast nucleic acids triggers a host-protective type I interferon response. Eur J Immunol 41: 1969–1979.

50. del Fresno, C., D. Soulat, S. Roth, K. Blazek, I. Udalova, D. Sancho, J. Ruland, and C. Ardavin. 2013. Interferon-beta production via Dectin-1-Syk-IRF5 signaling in dendritic cells is crucial for immunity to C. albicans. Immunity 38: 1176–1186.

51. Dutta, O., V. Espinosa, K. Wang, S. Avina, and A. Rivera. 2020. Dectin-1 Promotes Type I and III Interferon Expression to Support Optimal Antifungal Immunity in the Lung. Front Cell Infect Microbiol 10: 321.

52. Lin, J.D., N. Feng, A. Sen, M. Balan, H.C. Tseng, C. McElrath, S.V. Smirnov, J. Peng, L.L. Yasukawa, R.K. Durbin, J.E. Durbin, H.B. Greenberg, and S.V. Kotenko. 2016. Distinct Roles of Type I and Type III Interferons in Intestinal Immunity to Homologous and Heterologous Rotavirus Infections. PLoS Pathog 12: e1005600.

53. Hohl, T.M., H.L. Van Epps, A. Rivera, L.A. Morgan, P.L. Chen, M. Feldmesser, and E.G. Pamer. 2005. Aspergillus fumigatus triggers inflammatory responses by stage-specific beta-glucan display. PLoS Pathog 1: e30.

54. Szewczyk, E., T. Nayak, C.E. Oakley, H. Edgerton, Y. Xiong, N. Taheri-Talesh, S.A. Osmani, and B.R. Oakley. 2006. Fusion PCR and gene targeting in Aspergillus nidulans. Nat Protoc 1: 3111–3120.

55. Willger, S.D., S. Puttikamonkul, K.H. Kim, J.B. Burritt, N. Grahl, L.J. Metzler, R. Barbuch, M. Bard, C.B. Lawrence, and R.A. Cramer, Jr. 2008. A sterol-regulatory element binding protein is required for cell polarity, hypoxia adaptation, azole drug resistance, and virulence in Aspergillus fumigatus. PLoS Pathog 4: e1000200.

56. Jhingran, A., K.B. Mar, D.K. Kumasaka, S.E. Knoblaugh, L.Y. Ngo, B.H. Segal, Y. Iwakura, C.A. Lowell, J.A. Hamerman, X. Lin, and T.M. Hohl. 2012. Tracing conidial fate and measuring host cell antifungal activity using a reporter of microbial viability in the lung. Cell Rep 2: 1762–1773.

57. Schyns, J., F. Bureau, and T. Marichal. 2018. Lung Interstitial Macrophages: Past, Present, and Future. J Immunol Res 2018: 5160794.

58. Misharin, A.V., L. Morales-Nebreda, G.M. Mutlu, G.R. Budinger, and H. Perlman. 2013. Flow cytometric analysis of macrophages and dendritic cell subsets in the mouse lung. Am J Respir Cell Mol Biol 49: 503–510.

59. Steele, C., R.R. Rapaka, A. Metz, S.M. Pop, D.L. Williams, S. Gordon, J.K. Kolls, and G.D. Brown. 2005. The beta-glucan receptor dectin-1 recognizes specific morphologies of Aspergillus fumigatus. PLoS Pathog 1: e42.

60. Pichlmair, A., O. Schulz, C.P. Tan, J. Rehwinkel, H. Kato, O. Takeuchi, S. Akira, M. Way, G. Schiavo, and C. Reis e Sousa. 2009. Activation of MDA5 requires higher-order RNA structures generated during virus infection. J Virol 83: 10761–10769.

61. Pichlmair, A., O. Schulz, C.P. Tan, T.I. Naslund, P. Liljestrom, F. Weber, and C. Reis e Sousa. 2006. RIG-I-mediated antiviral responses to single-stranded RNA bearing 5’-phosphates. Science 314: 997–1001.

62. Odendall, C., E. Dixit, F. Stavru, H. Bierne, K.M. Franz, A.F. Durbin, S. Boulant, L. Gehrke, P. Cossart, and J.C. Kagan. 2014. Diverse intracellular pathogens activate type III interferon expression from peroxisomes. Nat Immunol 15: 717–726.

63. Jaeger, M., R. van der Lee, S.C. Cheng, M.D. Johnson, V. Kumar, A. Ng, T.S. Plantinga, S.P. Smeekens, M. Oosting, X. Wang, W. Barchet, K. Fitzgerald, L.A. Joosten, J.R. Perfect, C. Wijmenga, F.L. van de Veerdonk, M.A. Huynen, R.J. Xavier, B.J. Kullberg, and M.G. Netea. 2015. The RIG-I-like helicase receptor MDA5 (IFIH1) is involved in the host defense against Candida infections. Eur J Clin Microbiol Infect Dis 34: 963–974.

64. Groom, J.R. and A.D. Luster. 2011. CXCR3 ligands: redundant, collaborative and antagonistic functions. Immunol Cell Biol 89: 207–215.

65. Seth, R.B., L. Sun, C.K. Ea, and Z.J. Chen. 2005. Identification and characterization of MAVS, a mitochondrial antiviral signaling protein that activates NF-kappaB and IRF 3. Cell 122: 669–682.

66. Kawai, T., K. Takahashi, S. Sato, C. Coban, H. Kumar, H. Kato, K.J. Ishii, O. Takeuchi, and S. Akira. 2005. IPS-1, an adaptor triggering RIG-I- and Mda5-mediated type I interferon induction. Nat Immunol 6: 981–988.

67. Xu, L.G., Y.Y. Wang, K.J. Han, L.Y. Li, Z. Zhai, and H.B. Shu. 2005. VISA is an adapter protein required for virus-triggered IFN-beta signaling. Mol Cell 19: 727–740.

68. Meylan, E., J. Curran, K. Hofmann, D. Moradpour, M. Binder, R. Bartenschlager, and J. Tschopp. 2005. Cardif is an adaptor protein in the RIG-I antiviral pathway and is targeted by hepatitis C virus. Nature 437: 1167–1172.

69. Garlanda, C., E. Hirsch, S. Bozza, A. Salustri, M. De Acetis, R. Nota, A. Maccagno, F. Riva, B. Bottazzi, G. Peri, A. Doni, L. Vago, M. Botto, R. De Santis, P. Carminati, G. Siracusa, F. Altruda, A. Vecchi, L. Romani, and A. Mantovani. 2002. Non-redundant role of the long pentraxin PTX3 in anti-fungal innate immune response. Nature 420: 182–186.

70. Jhingran, A., S. Kasahara, K.M. Shepardson, B.A. Junecko, L.J. Heung, D.K. Kumasaka, S.E. Knoblaugh, X. Lin, B.I. Kazmierczak, T.A. Reinhart, R.A. Cramer, and T.M. Hohl. 2015. Compartment-specific and sequential role of MyD88 and CARD9 in chemokine induction and innate defense during respiratory fungal infection. PLoS Pathog 11: e1004589.

71. Drummond, R.A., A.L. Collar, M. Swamydas, C.A. Rodriguez, J.K. Lim, L.M. Mendez, D.L. Fink, A.P. Hsu, B. Zhai, H. Karauzum, C.M. Mikelis, S.R. Rose, E.M. Ferre, L. Yockey, K. Lemberg, H.S. Kuehn, S.D. Rosenzweig, X. Lin, P. Chittiboina, S.K. Datta, T.H. Belhorn, E.T. Weimer, M.L. Hernandez, T.M. Hohl, D.B. Kuhns, and M.S. Lionakis. 2015. CARD9-Dependent Neutrophil Recruitment Protects against Fungal Invasion of the Central Nervous System. PLoS Pathog 11: e1005293.

72. Kowalski, C.H., S.R. Beattie, K.K. Fuller, E.A. McGurk, Y.W. Tang, T.M. Hohl, J.J. Obar, and R.A. Cramer, Jr. 2016. Heterogeneity among Isolates Reveals that Fitness in Low Oxygen Correlates with Aspergillus fumigatus Virulence. mBio 7: :e01515–01516.

73. Fermaintt, C.S., K. Sano, Z. Liu, N. Ishii, J. Seino, N. Dobbs, T. Suzuki, Y.X. Fu, M.A. Lehrman, I. Matsuo, and N. Yan. 2019. A bioactive mammalian disaccharide associated with autoimmunity activates STING-TBK1-dependent immune response. Nat Commun 10: 2377.

74. Moretti, J., S. Roy, D. Bozec, J. Martinez, J.R. Chapman, B. Ueberheide, D.W. Lamming, Z.J. Chen, T. Horng, G. Yeretssian, D.R. Green, and J.M. Blander. 2017. STING Senses Microbial Viability to Orchestrate Stress-Mediated Autophagy of the Endoplasmic Reticulum. Cell 171: 809–823.

75. Alexopoulou, L., A.C. Holt, R. Medzhitov, and R.A. Flavell. 2001. Recognition of double-stranded RNA and activation of NF-kappaB by Toll-like receptor 3. Nature 413: 732–738.

76. Lund, J.M., L. Alexopoulou, A. Sato, M. Karow, N.C. Adams, N.W. Gale, A. Iwasaki, and R.A. Flavell. 2004. Recognition of single-stranded RNA viruses by Toll-like receptor 7. Proc Natl Acad Sci U S A 101: 5598–5603.

77. Gitlin, L., W. Barchet, S. Gilfillan, M. Cella, B. Beutler, R.A. Flavell, M.S. Diamond, and M. Colonna. 2006. Essential role of mda-5 in type I IFN responses to polyriboinosinic:polyribocytidylic acid and encephalomyocarditis picornavirus. Proc Natl Acad Sci U S A 103: 8459–8464.

78. Kato, H., O. Takeuchi, S. Sato, M. Yoneyama, M. Yamamoto, K. Matsui, S. Uematsu, A. Jung, T. Kawai, K.J. Ishii, O. Yamaguchi, K. Otsu, T. Tsujimura, C.S. Koh, C. Reis e Sousa, Y. Matsuura, T. Fujita, and S. Akira. 2006. Differential roles of MDA5 and RIG-I helicases in the recognition of RNA viruses. Nature 441: 101–105.

79. Yamamoto, M., S. Sato, K. Mori, K. Hoshino, O. Takeuchi, K. Takeda, and S. Akira. 2002. Cutting edge: a novel Toll/IL-1 receptor domain-containing adapter that preferentially activates the IFN-beta promoter in the Toll-like receptor signaling. J Immunol 169: 6668–6672.

80. Beisswenger, C., C. Hess, and R. Bals. 2012. Aspergillus fumigatus conidia induce interferon-beta signalling in respiratory epithelial cells. Eur Respir J 39: 411–418.

81. Davis, M.J. and J.A. Swanson. 2010. Technical advance: Caspase-1 activation and IL-1beta release correlate with the degree of lysosome damage, as illustrated by a novel imaging method to quantify phagolysosome damage. J Leukoc Biol 88: 813–822.

82. Hornung, V., F. Bauernfeind, A. Halle, E.O. Samstad, H. Kono, K.L. Rock, K.A. Fitzgerald, and E. Latz. 2008. Silica crystals and aluminum salts activate the NALP3 inflammasome through phagosomal destabilization. Nat Immunol 9: 847–856.

83. Wellington, M., K. Koselny, and D.J. Krysan. 2012. Candida albicans morphogenesis is not required for macrophage interleukin 1beta production. mBio 4: e00433–00412.

84. Nguyen, T.A., B.R.C. Smith, M.D. Tate, G.T. Belz, M.H. Barrios, K.D. Elgass, A.S. Weisman, P.J. Baker, S.P. Preston, L. Whitehead, A. Garnham, R.J. Lundie, G.K. Smyth, M. Pellegrini, M. O’Keeffe, I.P. Wicks, S.L. Masters, C.P. Hunter, and K.C. Pang. 2017. SIDT2 Transports Extracellular dsRNA into the Cytoplasm for Innate Immune Recognition. Immunity 47: 498–509 e496.

85. Joffe, L.S., L. Nimrichter, M.L. Rodrigues, and M. Del Poeta. 2016. Potential Roles of Fungal Extracellular Vesicles during Infection. mSphere 1:

86. Cheng, Y. and J.S. Schorey. 2019. Extracellular vesicles deliver Mycobacterium RNA to promote host immunity and bacterial killing. EMBO Rep 20:

87. Singh, P.P., L. Li, and J.S. Schorey. 2015. Exosomal RNA from Mycobacterium tuberculosis-Infected Cells Is Functional in Recipient Macrophages. Traffic 16: 555–571.

88. Mehrad, B., R.M. Strieter, T.A. Moore, W.C. Tsai, S.A. Lira, and T.J. Standiford. 1999. CXC chemokine receptor-2 ligands are necessary components of neutrophil-mediated host defense in invasive pulmonary aspergillosis. J Immunol 163: 6086–6094.

89. Lang, S., L. Li, X. Wang, J. Sun, X. Xue, Y. Xiao, M. Zhang, T. Ao, and J. Wang. 2017. CXCL10/IP-10 Neutralization Can Ameliorate Lipopolysaccharide-Induced Acute Respiratory Distress Syndrome in Rats. PLoS One 12: e0169100.

90. Ichikawa, A., K. Kuba, M. Morita, S. Chida, H. Tezuka, H. Hara, T. Sasaki, T. Ohteki, V.M. Ranieri, C.C. dos Santos, Y. Kawaoka, S. Akira, A.D. Luster, B. Lu, J.M. Penninger, S. Uhlig, A.S. Slutsky, and Y. Imai. 2013. CXCL10-CXCR3 enhances the development of neutrophil-mediated fulminant lung injury of viral and nonviral origin. Am J Respir Crit Care Med 187: 65–77.

91. Muller, U., U. Steinhoff, L.F. Reis, S. Hemmi, J. Pavlovic, R.M. Zinkernagel, and M. Aguet. 1994. Functional role of type I and type II interferons in antiviral defense. Science 264: 1918–1921.

92. Ng, C.T., B.M. Sullivan, J.R. Teijaro, A.M. Lee, M. Welch, S. Rice, K.C. Sheehan, R.D. Schreiber, and M.B. Oldstone. 2015. Blockade of interferon Beta, but not interferon alpha, signaling controls persistent viral infection. Cell Host Microbe 17: 653–661.

93. Teijaro, J.R., C. Ng, A.M. Lee, B.M. Sullivan, K.C. Sheehan, M. Welch, R.D. Schreiber, J.C. de la Torre, and M.B. Oldstone. 2013. Persistent LCMV infection is controlled by blockade of type I interferon signaling. Science 340: 207–211.

94. Wilson, E.B., D.H. Yamada, H. Elsaesser, J. Herskovitz, J. Deng, G. Cheng, B.J. Aronow, C.L. Karp, and D.G. Brooks. 2013. Blockade of chronic type I interferon signaling to control persistent LCMV infection. Science 340: 202–207.

95. Seyedmousavi, S., M.J. Davis, J.A. Sugui, T. Pinkhasov, S. Moyer, A.M. Salazar, Y.C. Chang, and K.J. Kwon-Chung. 2018. Exogenous Stimulation of Type I Interferon Protects Mice with Chronic Granulomatous Disease from Aspergillosis through Early Recruitment of Host-Protective Neutrophils into the Lung. mBio 9:

96. Sionov, E., K.D. Mayer-Barber, Y.C. Chang, K.D. Kauffman, M.A. Eckhaus, A.M. Salazar, D.L. Barber, and K.J. Kwon-Chung. 2015. Type I IFN Induction via Poly-ICLC Protects Mice against Cryptococcosis. PLoS Pathog 11: e1005040.

97. Davis, M.J., S. Moyer, E.S. Hoke, E. Sionov, K.D. Mayer-Barber, D.L. Barber, H. Cai, L. Jenkins, P.J. Walter, Y.C. Chang, and K.J. Kwon-Chung. 2019. Pulmonary Iron Limitation Induced by Exogenous Type I IFN Protects Mice from Cryptococcus gattii Independently of T Cells. mBio 10:

98. Biondo, C., A. Midiri, M. Gambuzza, E. Gerace, M. Falduto, R. Galbo, A. Bellantoni, C. Beninati, G. Teti, T. Leanderson, and G. Mancuso. 2008. IFN-alpha/beta signaling is required for polarization of cytokine responses toward a protective type 1 pattern during experimental cryptococcosis. J Immunol 181: 566–573.

99. Sato, K., H. Yamamoto, T. Nomura, I. Matsumoto, T. Miyasaka, T. Zong, E. Kanno, K. Uno, K. Ishii, and K. Kawakami. 2015. Cryptococcus neoformans Infection in Mice Lacking Type I Interferon Signaling Leads to Increased Fungal Clearance and IL-4-Dependent Mucin Production in the Lungs. PLoS One 10: e0138291.

100. Van Prooyen, N., C.A. Henderson, D. Hocking Murray, and A. Sil. 2016. CD103+ Conventional Dendritic Cells Are Critical for TLR7/9-Dependent Host Defense against Histoplasma capsulatum, an Endemic Fungal Pathogen of Humans. PLoS Pathog 12: e1005749.

101. Riedelberger, M., P. Penninger, M. Tscherner, M. Seifert, S. Jenull, C. Brunnhofer, B. Scheidl, I. Tsymala, C. Bourgeois, A. Petryshyn, W. Glaser, A. Limbeck, B. Strobl, G. Weiss, and K. Kuchler. 2020. Type I Interferon Response Dysregulates Host Iron Homeostasis and Enhances Candida glabrata Infection. Cell Host Microbe 27: 454–466 e458.

102. Riedelberger, M., P. Penninger, M. Tscherner, B. Hadriga, C. Brunnhofer, S. Jenull, A. Stoiber, C. Bourgeois, A. Petryshyn, W. Glaser, A. Limbeck, M.A. Lynes, G. Schabbauer, G. Weiss, and K. Kuchler. 2020. Type I Interferons Ameliorate Zinc Intoxication of Candida glabrata by Macrophages and Promote Fungal Immune Evasion. iScience 23: 101121.

103. Bourgeois, C., O. Majer, I.E. Frohner, I. Lesiak-Markowicz, K.S. Hildering, W. Glaser, S. Stockinger, T. Decker, S. Akira, M. Muller, and K. Kuchler. 2011. Conventional dendritic cells mount a type I IFN response against Candida spp. requiring novel phagosomal TLR7-mediated IFN-beta signaling. J Immunol 186: 3104–3112.

104. Smeekens, S.P., A. Ng, V. Kumar, M.D. Johnson, T.S. Plantinga, C. van Diemen, P. Arts, E.T. Verwiel, M.S. Gresnigt, K. Fransen, S. van Sommeren, M. Oosting, S.C. Cheng, L.A. Joosten, A. Hoischen, B.J. Kullberg, W.K. Scott, J.R. Perfect, J.W. van der Meer, C. Wijmenga, M.G. Netea, and R.J. Xavier. 2013. Functional genomics identifies type I interferon pathway as central for host defense against Candida albicans. Nat Commun 4: 1342.

105. Majer, O., C. Bourgeois, F. Zwolanek, C. Lassnig, D. Kerjaschki, M. Mack, M. Muller, and K. Kuchler. 2012. Type I interferons promote fatal immunopathology by regulating inflammatory monocytes and neutrophils during Candida infections. PLoS Pathog 8: e1002811.

106. Hagmann, C.A., A.M. Herzner, Z. Abdullah, T. Zillinger, C. Jakobs, C. Schuberth, C. Coch, P.G. Higgins, H. Wisplinghoff, W. Barchet, V. Hornung, G. Hartmann, and M. Schlee. 2013. RIG-I detects triphosphorylated RNA of Listeria monocytogenes during infection in non-immune cells. PLoS One 8: e62872.

107. Abdullah, Z., M. Schlee, S. Roth, M.A. Mraheil, W. Barchet, J. Bottcher, T. Hain, S. Geiger, Y. Hayakawa, J.H. Fritz, F. Civril, K.P. Hopfner, C. Kurts, J. Ruland, G. Hartmann, T. Chakraborty, and P.A. Knolle. 2012. RIG-I detects infection with live Listeria by sensing secreted bacterial nucleic acids. EMBO J 31: 4153–4164.

108. Liehl, P., V. Zuzarte-Luis, J. Chan, T. Zillinger, F. Baptista, D. Carapau, M. Konert, K.K. Hanson, C. Carret, C. Lassnig, M. Muller, U. Kalinke, M. Saeed, A.F. Chora, D.T. Golenbock, B. Strobl, M. Prudencio, L.P. Coelho, S.H. Kappe, G. Superti-Furga, A. Pichlmair, A.M. Vigario, C.M. Rice, K.A. Fitzgerald, W. Barchet, and M.M. Mota. 2014. Host-cell sensors for Plasmodium activate innate immunity against liver-stage infection. Nat Med 20: 47–53.

109. Cheng, Y. and J.S. Schorey. 2018. Mycobacterium tuberculosis-induced IFN-beta production requires cytosolic DNA and RNA sensing pathways. J Exp Med 215: 2919–2935.

