## Supplement Figures for "MDA5 is an essential vita-PAMP sensor necessary for host resistance against *Aspergillus fumigatus*"

**Supplemental Table 1. Primers used to construct the CEA10 FLARE strain.**

| Primer name | Sequence (5' → 3') |
|---|---|
| **RAC2888** | agagtatgcggcaagtcatgagcagttccaggtggaatgttatg |
| **RAC2799** | Gtgatgtctgctcaagcggggta |
| **RAC4582** | taccccgcttgagcagacatcacatgactggcggcaaatctggtgg |
| **RAC4583** | ggctccagcgcctgcaccagctcccagctcctggctgccctttccg |
| **RAC2600** | Ggagctggtgcaggcgctggagcc |
| **RAC4575** | gcgttttattcttgttgacatgggttaggcgccggtggagtggcggc |
| **RAC2536** | Cccatgtcaacaagaataaaacgc |
| **RAC2537** | Ccgagtggagatgtggagt |
| **RAC1981** | gagcaattggaatccgcccctccgccccgagctcccaaatctgtccagatc |
| **RAC4134** | agagcattgtttgaggcgaccggttacctgtgcattctgggtaaacgactc |
| **RAC3873** | gcagctaactctggcagcacc |
| **RAC3874** | ggcggaggggcggattccaattgctccgtcacttctcatgctacggacac |
| **RAC3875** | cgcatcagtgcctcctctcagacgagtgaggcttggggttcgtc |
| **RAC3876** | gttctgcccctggatcgctagg |
| **RAC2055** | accggtcgcctcaaacaatgctct |
| **RAC2056** | gtctgagaggaggcactgatgcg |
| **RAC3877** | gacgccggttctctgggtaatc |
| **RAC3878** | cacgggataaccatgcaataggcg |

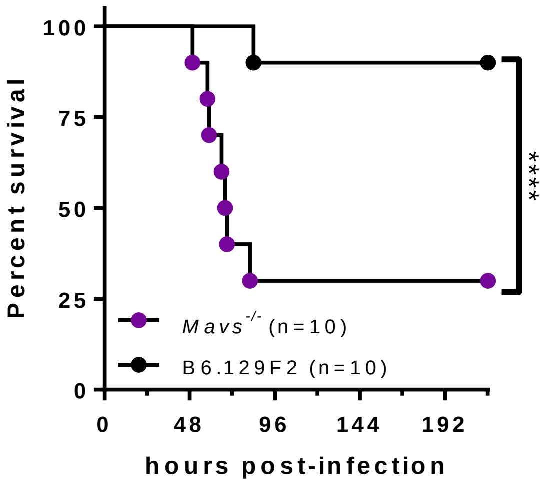


__Supplemental Figure 1.__ ***Mavs*-dependent responses are essential for host resistance against the Af293 strain of Aspergillus fumigatus.** *Mavs*$^{-/-}$ and B6.129F2 mice were challenged i.t. with $4x10^7$ resting conidia of the Af293 isolate of *Aspergillus fumigatus.* Survival analysis in immune-competent wild-type and knock-out mice were tracked over the first 9 days. ****, P < 0.0001 by Mantel-Cox log rank test.

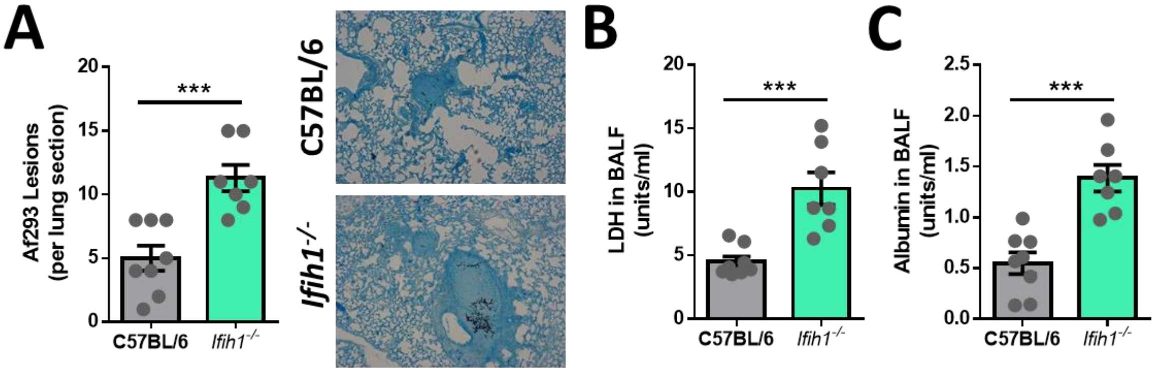


<u>Supplemental Figure 2.</u> **Mda5-dependent responses are essential for host resistance against invasive aspergillosis in corticosteroid immunosuppressed mice.** Mice were treated with 40 mg/kg of Kenalog given s.c. on one day prior to and 4 days after challenge i.t. with $10^6$ resting conidia of the Af293 strain of *A. fumigatus.* At 72 h post-inoculation lungs were collected for histological analysis for fungal growth **(A).** BALF was also collected at 72 h post-inoculation to assess LDH activity as a proxy for lung damage **(B)** and albumin as a proxy for lung leakage **(C)**. Mann-Whitney U-test was used for statistical analysis.

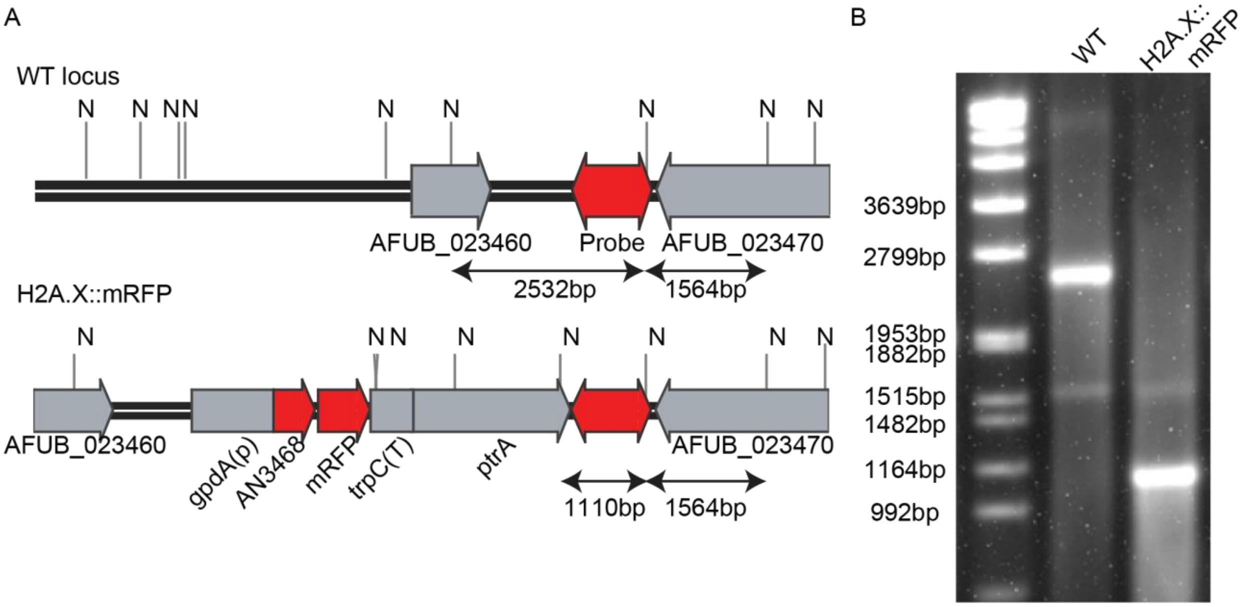


**<u>Supplemental Figure 3. DNA analysis for confirmation of CEA10 H2A.X::mRFP strain.</u> (A)** Schematic representation showing the intergenic distance between AFUB_023460 and AFUB_023470. H2A.X (AN3468) from *A. nidulans* was fused with *gpdA*(P), *trpC*(T) and *ptrA* gene conferring resistance to pyrithiamine as described in materials and methods. The DNA fragment used as probe and the expected DNA hybridization patterns for southern blot analysis are shown. **(B)** Southern blot analysis of CEA10 WT and H2AX:mRFP strain. NsiI digested DNA of each strain was hybridized with the probe as shown in A. N – *Nsi*I restriction site.
